# Competitive specific anchorage of molecules onto surfaces: quantitative control of grafting densities and contamination by free anchors

**DOI:** 10.1101/2023.06.27.546772

**Authors:** Oksana Kirichuk, Sumitra Srimasorn, Xiaoli Zhang, Abigail R. E. Roberts, Liliane Coche-Guerente, Jessica C. F. Kwok, Lionel Bureau, Delphine Débarre, Ralf P. Richter

**Affiliations:** School of Biomedical Sciences, Faculty of Biological Sciences, University of Leeds, Leeds LS2 9JT, United Kingdom; School of Physics and Astronomy, Faculty of Engineering and Physical Sciences, Astbury Centre for Structural Molecular Biology, and Bragg Centre for Materials Research, University of Leeds, Leeds LS2 9JT, United Kingdom; Université Grenoble-Alpes, CNRS, LIPhy, 38000 Grenoble, France; Université Grenoble-Alpes, CNRS, Département de Chimie Moléculaire, 38000 Grenoble, France; Institute of Experimental Medicine, Czech Academy of Sciences, Vídeňská, 1083 Prague, Czech Republic

**Author notes:** Corresponding authors: D.D. -; R.P.R. –.

**Keywords:** streptavidin, biotin, quartz crystal microbalance, ellipsometry, fluorescence, surface density

## Abstract

The formation of surfaces decorated with biomacromolecules such as proteins, glycans or nucleic acids with well-controlled orientations and densities is of critical importance for the design of *in vitro* models, *e.g*., synthetic cell membranes, and interaction assays. To this effect, ligand molecules are often functionalized with an anchor that specifically binds to a surface with a high density of binding sites, providing control over the presentation of the molecules. Here, we present a method to robustly and quantitatively control the surface density of one or several types of anchor-bearing molecules by tuning the relative concentrations of target molecules and free anchors in the incubation solution. We provide a theoretical background that relates incubation concentrations to the final surface density of the molecules of interest, and present effective guidelines towards optimizing incubation conditions for the quantitative control of surface densities. Focussing on the biotin anchor, a commonly used anchor for interaction studies, as a salient example, we experimentally demonstrate surface density control over a wide range of densities and target molecule sizes. Conversely, we show how the method can be adapted to quality control the purity of end-grafted biopolymers such as biotinylated glycosaminoglycans by quantifying the amount of residual free biotin reactant in the sample solution.

## INTRODUCTION

Tuning the density of surface-anchored biomacromolecules, such as proteins, glycans or nucleic acids, is desirable in a wide range of applications. In biomolecular interaction assays, for example, the binding of multivalent analytes to surface-anchored ligands depends sensitively on the ligand surface density, and quantitative tuning of the ligand surface density enables avidity effects to be probed ^1-2^. Similarly, the response of cells to model surfaces presenting ligands for binding to cognate cell surface receptors often depends sensitively on the ligand surface density, and ligand density tuning thus can differentially impact downstream intracellular signalling and cell phenotype such as adhesion, migration and differentiation ^3-6^. Proper control over ligand surface densities is also beneficial to maximize the selectivity and yield of separation, for example, in affinity chromatography or bead-based pathogen or cell capture devices. A case in point is the emerging concept of ‘superselective’ binding, which entirely relies on the sharp discrimination of surfaces by their ligand density ^7-8^.

Despite the established need, it remains far from trivial to coat surfaces with biomacromolecular ligands at quantitatively tuneable densities. Crucially, a suitable passivation of the surface and a control of ligand orientation are required alongside the control of ligand density, to impede non-specific interactions with the surface and to retain the functionality of surface bound biomacromolecules. Most methods control the level of ligand binding to the surface from a solution of ligands, as reviewed in Ref. 2. A number of techniques couple passivation and functionalization with ligands through the formation of a mixed self-assembled monolayer (SAM ^3, 9-11^) or a supported lipid bilayer (SLB ^12-15^) by incubation with a mix of inert and active molecules at a defined ratio. The active molecule can directly couple the ligand of interest ^15-16^, or a reactive group or intermediate ligand allowing subsequent coupling after the layer formation ^17-18^. Often, however, the relation between the ligand density on the surface and the initial incubation mix is not a straightforward one, although subsequent quantification is possible ^11, 18-22^. An alternative is to functionalize modified polymers such as PLL-*g*-PEG or DNA with a controlled density of ligands that translates into a chosen surface density upon adsorption onto a surface ^23-25^. In this approach, however, the ligand of interest is not segregated from the macromolecule cover and the underlying surface, and hence access to the ligand may be sterically hindered and heterogeneous ^25-26^.

To achieve better control over the density of properly-exposed functional ligands, other methods rely on a passivating platform that is subsequently functionalized, such as biotinylated SLBs or SAMs covered with a well-defined surface density of stably bound streptavidin ^27-28^. Indeed, a number of methods have been developed that rely on the chemical conjugation of an anchor moiety at a defined (and suitable) site of the biomacromolecule of interest along with surfaces that are conceived to bind (or more accurately ‘graft’) the anchors with high affinity (for stable attachment) and high specificity (for selective attachment). One of the most popular anchor tags is biotin, which binds biotin binding proteins such as streptavidin. The high affinity of the biotin-streptavidin interaction (K_d_ ≈ 10^−14^ M ^29^) and the relative simplicity of biomacromolecule biotinylation ^30-31^ have resulted in a wide range of applications of this model system ^27-28, 32^.

The surface density of biotinylated ligands can then be controlled through different approaches. One common method relies on binding kinetics: ligand concentration and/or incubation time are tuned to achieve the desired ligand coverage ^33-36^. This approach is relatively simple but has its drawbacks. Firstly, the incubation time and ligand concentration need to be tightly controlled, as they are sensitively (typically linearly) affecting the final ligand surface density. Secondly, binding is often limited (in part or in full) by the diffusive and/or convective transport of ligands to the surface, which makes the binding rate sensitive to the specific incubation conditions. When samples are incubated in stagnant solutions, for example, they typically require an initial phase of convective mixing for solution homogenization and a final phase of convective mixing to remove excess ligands from the solution phase. Binding throughout these transient phases can make a substantial contribution to the binding process ^37^, thus adding errors to the ligand surface density. Moreover, the mass-transport conditions in a specific fluidic device are often not accurately known, as they depend sensitively on factors such as temperature, solution viscosity, flow geometry and rate. This makes quantitative predictions of binding rates difficult and thus requires experimental calibration. It also reduces reproducibility as incubation conditions that have been established for one device cannot be readily transferred to another device (*e*.*g*., with a different flow geometry).

Another approach is to rely on ligand depletion to control the ligand surface density ^38^: the incubation time is chosen long enough to ensure adsorption of all molecules from the solution and the method is thus insensitive to mass transport conditions and the exact incubation time. However, the depletion method is (linearly) sensitive to the initial ligand concentration in solution, and this sensitivity is exacerbated by the small concentration values that are typically required to functionalize a surface at densities smaller than saturation, resulting in a significant influence of non-specific adsorption onto surfaces other than the one to functionalize.

Finally, the final density of grafted ligand can be tuned by mixing the biotinylated molecule of interest with other biotinylated molecules of similar size and chemical properties ^20^-^21, 39^. In this scenario, the final ligand surface density becomes insensitive to the incubation time (as long as surface saturation is attained) and absolute ligand concentration. Instead, it is mainly controlled by the mixing ratio, with the common assumption that the mixing ratio on the surface is equal to the mixing ratio in solution. This assumption, however, only holds for molecules of similar size and binding properties, which may require specific synthesis to achieve, thereby limiting the application and ease of use of this method.

In the present work, we build on this empirical approach to demonstrate a generic, versatile method to control the density of one or more ligands by competitive adsorption of the anchor-tagged ligand(s) and the free anchor itself. Our method relies on the fact that many anchors are rapid binders, so that mass-transport limited binding conditions are easily achieved: in these conditions, ligand grafting density can be quantitatively controlled by adapting the mixing ratio in solution to the ratio of hydrodynamic radii of the competing species. We focus on streptavidin-coated surfaces and biotin anchors to validate our method experimentally. However, the approach should be equally suitable for any other tag with a sufficiently high intrinsic binding rate, such as nickel chelating surfaces to capture polyhistidine tags on proteins ^40-42^, surfaces coated with protein A, protein G or their functional parts to capture antibodies or fusion proteins with an Fc tag ^28^, and DNA coated surfaces to capture specific sections of mRNA or DNA strands.

Importantly, we provide the theoretical background to relate solution and surface fractions of ligands in different incubation conditions, and a set of guidelines to facilitate quantitative tuning of the ligand grafting density. Moreover, we illustrate that the competitive binding concept can be expanded to purposes other than tuning the ligand surface density by demonstrating how it can be deployed to quantify the contamination with free anchors of complex biomacromolecules that are difficult to analyse with conventional chemical methods.

## MATERIALS AND METHODS

### Materials

Lyophilized 1,2-dioleoyl-sn-glycero-3-phosphocholine (DOPC) and 1,2-dioleoyl-sn-glycero-3-phosphoethanolamine-N-(Cap Biotinyl) (DOPE-cap-B) were purchased from Avanti Polar Lipids (Alabaster, USA). Lyophilized streptavidin (SAv; ∼60 kDa) was purchased from Sigma-Aldrich (# S4762).

Biotin (244.3 Da) was purchased from Sigma-Aldrich (# B4639). Biotinylated fluoresceine isothiocyanate (b-FITC; 732.8 Da) was purchased from Thermo Scientific (# 10752905). A tandem repeat of the Z domain of protein A connected through a flexible spacer (12 amino acids) to an N-terminal biotin (b-ZZ; 16.2 kDa) was expressed in *E. coli* and purified as described in detail elsewhere ^27^.

A recombinant P-selectin-Fc fusion protein (R&D Systems # 137-PS; ∼300 kDa) was purchased from Bio-Techne (Abingdon, UK). The construct contained at its N-terminus the amino acids Trp42 to Ala771 of human P-selectin, representing the receptor’s ectodomain, followed by a spacer of 7 amino acids, and the amino acids Pro100 to Lys330 of human IgG1. The homodimer construct was glycosylated, with a molecular mass estimated by the manufacturer to lie between 146 and 160 kDa per protomer according to SDS polyacrylamide gel electrophoresis.

Chondroitin sulphate glycosaminoglycan CS-D, derived from Shark cartilage and with a mean molecular mass of 30 kDa, was purchased from AMSBIO (Abingdon, UK; # 400676). CS-D is an unbranched polysaccharide consisting of β(1,4)-glucuronic acid (GlcA)-β(1,3)-N-acetyl galactosamine (GlcNAc) disaccharides, where the C2 position in GlcA and the C6 position in GlcNAc are preferentially sulfated.

HEPES buffered saline (HBS; 10 mM HEPES, pH 7.4, 150 mM NaCl) was prepared in ultrapure water (resistivity 18.2 MΩ·cm) and used as working buffer throughout all the measurements.

Small unilamellar vesicles (SUVs) containing 5 mol-% biotin were prepared as previously described ^43^. Briefly, lipids were dissolved in chloroform, mixed at a molar ratio of 95 % DOPC and 5 % DOPE-cap-B, dried under a stream of nitrogen gas followed by drying in a vacuum desiccator for at least 2 h. The lipid mixture was then resuspended in HBS at a final concentration of 2 mg/ml and homogenized by five cycles of freezing, thawing and vortexing. The lipid suspension was sonicated with a tip sonicator (FB120; Fisher Scientific, UK) in pulsed mode (duty cycle: 1 s on (at 70% maximal power) / 1 s off) with refrigeration for a total time of 30 min, followed by centrifugation (10 min at 12,100 × *g*) to remove titanium particle debris from the sonicator tip. SUVs were stored at 4 °C under an inert gas (nitrogen or argon) until use.

### Quartz crystal microbalance with dissipation monitoring (QCM-D)

QCM-D measurements were performed with a Q-Sense E4 system equipped with Flow Modules (Biolin Scientific, Västra Frölunda, Sweden) on silica-coated sensors (QSX303; Biolin Scientific). The flow rate was controlled with a syringe pump (Legato; World Precision Instruments, Stevenage, UK) at 20 μl/min unless otherwise stated. The working temperature was typically 23 °C, except for application example 3 where it was 25 °C. Before each use, sensors were cleaned in a 2% (*w*/*v*) aqueous solution of sodium dodecyl sulfate (SDS) detergent for 30 min, rinsed with ultrapure water, blow dried with N2 gas, and treated with UV/ozone for another 30 min. QCM-D data were collected at six overtones (*i* = 3, 5, 7, 9, 11 and 13, corresponding to resonance frequencies of approximately 15, 25, 35, 45, 55, and 65 MHz). Changes in normalized resonance frequency (Δ*F* = Δ*fi*/*i*) and dissipation (Δ*D*) of the fifth overtone (*i* = 5) are presented. All other overtones provided comparable information.

The thickness of dense protein monolayers was estimated from the QCM-D frequency shift using the Sauerbrey equation, as *h* = −*C*Δ*F⁄ρ*, with the mass-sensitivity constant *C* = 18.0 ng/(cm^2^ Hz). The film density was assumed to be 1.1 g/cm3, reflecting the solvated nature of the film to a good approximation. With a typical volume density of 1.36 g/cm3 for proteins in aqueous solvent ^44^, and a density of 1.0 g/cm3 for water, the effective film density of 1.1 g/cm3 corresponds to a 1:2 mass ratio of protein and solvent. With this assumption, film thickness errors owing to incorrect film density estimates are inferior to 10% for protein-to-solvent mass ratios up to 2:1, which covers even very dense (*e*.*g*., crystalline) protein layers ^45^. We verified that Δ*D⁄*−Δ*F* ≪ 0.4 ppm*⁄*Hz (and that the Δ*F* curves essentially overlay across all overtones), to ascertain films are sufficiently rigid for the Sauerbrey equation to provide reliable film thickness estimates ^46^.

### Spectroscopic ellipsometry (SE)

SE measures changes in the polarisation of light upon reflection at a planar surface. SE measurements were performed *in situ* in a custom-built open cuvette (∼100 μl volume) with glass windows, on silicon wafers, at room temperature with a spectroscopic rotating compensator ellipsometer with a horizontal plane of incidence (M-2000 V; J. A. Woollam, Lincoln, NE). Ellipsometric angles Δ and Ψ were acquired over a wavelength range from λ = 370 to 1000 nm, at an angle of incidence of 70° and with a time resolution of 5 to 10 s. All samples (in working buffer) were directly pipetted into the cuvette, and homogenized by a magnetic stirrer, located at the bottom of the cuvette. SUVs were incubated under continuous stirring. All other samples were stirred for approximately 5 s after sample injection, and for the remainder of the sample incubation time, the stirrer was turned off and adsorption was left to proceed from stagnant solution. Excess sample was rinsed away by flowing working buffer through the cuvette; this was assisted by a flow-through tubing system and a peristaltic pump (IPC; Ismatec, Germany) operated at a flow rate of 5 ml/min; during the rinsing phases, the stirrer was turned on to ensure homogenization and maximize exchange of the cuvette content.

Areal mass densities (*AMD*) and molar surface densities of adsorbed biomolecules were determined by numerical fitting of the SE data using the software CompleteEASE (J. A. Woollam). A model with a stack of multiple isotropic layers relates the measured ellipsometric angles Δ and Ψ as a function of λ to the optical properties of the substrate, the adsorbed films and the surrounding buffer solution. The semi-infinite bulk solution was treated as a transparent Cauchy medium (refractive index: *n*sol(λ) = *A*_sol_ + *B*_sol_/λ^2^, where *A*_sol_ = 1.325 and *B*_sol_ = 0.00322 μm^2 47^). The native oxide film on the Si wafers was modelled as a single and transparent Cauchy layer. Its optical properties were determined from the measurements acquired in the presence of bulk solution but in the absence of the biomolecular film, which were then fitted over the range of λ using the tabulated values for the underlying Si substrate (implemented in CompleteEASE) ^27^. The adsorbed biomolecular film was fitted with the help of two separate layers. The combination of SLB, the monolayer of streptavidin and b-ZZ adapter protein (or the mix of b-ZZ with biotin) was treated as a single layer (index 1), which was treated as a transparent Cauchy medium with thickness (*d*1) and a wavelength-dependent refractive index *n*1(λ) = *A*1 + *B*1/λ^2^. As this layer was thin (*d*_1_ < 10 nm), the optical parameter *A*1 and *B*1 were kept fixed, and *d*1 was the only adjustable parameter. A1 = 1.4 was set as a typical value for a solvated biomolecular film, and the dispersity was set equal to the bulk solution (*B*1 = *B*sol). The thicker layer of P-selectin, adsorbed on b-ZZ, was treated as a separate transparent Cauchy layer (index 2) with thickness *d*_2_ and *n*_2_(λ) = *A*_2_ + *B*_2_/λ^2^. Here, *d*_2_ and *A*_2_ were adjustable fitting parameters, and the change in *B*_2_ with the protein concentration was neglected so that *B*_2_ = *B*_sol_. Layer 1 was assumed to remain unchanged during P-selectin binding. The root mean square error remained typically below 2 throughout the time-resolved data fitting, which indicated a good fit. The areal mass densities (*AMD*) were determined through a variant of de Fejter’s equation ^48^, *AMD* = [*d*(*A*− *A*_sol_)]*⁄*(d*n⁄*d*c*) using the refractive index increments, d*n*/d*c*, of 0.18 cm3/g for all proteins, and 0.169 cm^3^/g for lipids. The molar surface density was calculated from the areal mass density for b-ZZ and P-selectin as Γ = *AMD*/*M*_w_, where *M*_w_ is the molecular weight of the protein. Errors in *AMD* and Γ comprise the temporal noise and the confidence intervals of the data fitting.

### Surface preparation for confocal microscopy

Confocal microscopy analysis was performed on glass coverslips of 35 mm diameter (VWR, USA). The coverslips were cleaned with Piranha solution (H_2_O_2_:H_2_SO_4_ = 1:3), and exposed to H_2_O plasma (Plasma surface cleaning system; Diener Electronic, Germany) for 3 min immediately before use. Each coverslip was then mounted on a custom-made Teflon holder with the help of a two-component glue (Twinsil, Picodent, Germany) so that the coverslip formed the bottom of four identical wells. The cylindrical wells had a diameter of 5 mm and a volume of 50 μl.

Supported lipid bilayers (SLBs) were formed by the method of vesicle rupture and spreading. Surfaces were incubated with 50 μg/ml SUVs in working buffer for 30 min allowing the SLB to form. Excess SUVs were removed by 10 washes with working buffer (in each wash, 100 μl HBS were injected into the well, the solution was mixed with a pipette taking care not to touch the bottom of the well, and 100 μl of liquid were removed). The SLB-coated surface was then incubated with 0.33 μM (20 μg/ml) SAv in working buffer for 60 min, to form a dense SAv monolayer presenting biotin-binding sites, and excess SAv was again removed by washing 10 times with working buffer as described above.

For further functionalisation with b-FITC, the original fluorophore solution in working buffer was deliberately partially photobleached, to reduce the concentration of fluorescent b-FITC to a level that essentially avoided self-quenching on the surface (for details, see Figure S1). The resulting b-FITC solution (with a defined total concentration of fluorescent and non-fluorescent molecules) was then mixed with free biotin at the desired ratio. SAv-coated surfaces were incubated with biotinylated molecules (14 μM total concentration) in working buffer for 14 h, and washed 10 times with working buffer. Such a long incubation time was used, as it was found to improve the homogeneity of the layer when inspected by fluorescence microscopy.

### Confocal microscopy and image analysis

The fluorescence intensity of the functionalized surfaces was measured using a confocal laser scanning microscope (TCS SP8, Leica, Germany) using a 40×/1.30 oil objective and a built-in autofocus. Fluorescence was excited at 488 nm with a power on the sample in the range 0.5-10 μW, and detected in the wavelength range 491-629 nm with a pixel dwell time of 1.2 μs and a sampling of 0.284 μm/pixel. The surface coating was mostly uniform, as assessed by the fluorescence intensity distribution, although minor tilt of the sample resulted in gradients of intensity across the image. The autofocus was set such that the maximum intensity (corresponding to an in-focus surface) was located around the centre of the image, ensuring that the in-focus signal was reliably detected in all images. A mosaic of 40 to 100 images, each 292.6 μm × 292.6 μm in size, was then acquired, so as to sample the entire surface of each well.

Acquired images were analyzed with Fiji using custom routines. A band of 200 pixels in width was drawn across the image such that it cut through the region of maximal intensity, and the profile of the mean grey values across the band was defined. The profile was fitted with a polynomial function and the maximum value of the fit function in the field of view was considered as the in-focus intensity of a given image. Intensity values represent the mean ± standard error of the in-focus intensity across all images in a mosaic.

### Biotinylation of GAGs, and separation of GAGs from free biotin

CS-D GAG polysaccharides were biotinylated following the protocol described by Thakar *et al*. ^49^, with modifications. As the biotin derivative, we used alkoxyamine-EG_4_-biotin (Thermofisher Scientific # 26137; 434.2 Da), resulting in a =N-EG_4_-NH-CO-(CH_2_)_4_-biotin moiety at the C1 of the GAG’s reducing-end. Reactants were incubated at final concentrations of 1.2 mg/ml GAG, 20 mM aniline (Sigma Aldrich) and 150 μM alkoxyamine-EG4-biotin in 50 mM acetate buffer, pH 4.5. The reaction volume was typically 0.2 ml, and the mixture was left to react at 37 °C shaking at 300 rpm overnight.

GAGs were purified with a column of 7 mm diameter and 10 cm length (BioRad) packed with Sephadex G25 resin (Sigma-Aldrich # G25150). Dry resin was allowed to swell at room temperature for 3 h in 25% (*v*/*v*) ethanol solution, and then to settle before removing the supernatant. A slurry of 75% resin and 25% ultrapure water (*v*/*v*) was then prepared, degassed, and added to the chromatography column, taking care not to introduce bubbles. The resin was allowed to settle by gravity, the flow through was discarded, and the packed column was equilibrated with ultrapure water (3× 4 ml) before sample addition.

For purification, the 0.2 ml sample mixture was added to, and allowed to enter, the column’s resin bed. 0.7 ml of ultrapure water were then added and allowed to enter the bed, and the flow through was discarded. 2 ml of ultrapure water were then added, and the eluate was collected in fractions of 250 μl. The collected fractions were further characterized by QCM-D, or stored at -20 °C until use.

The average size of the GAGs (in number of disaccharides, *n*_ds_, of the linear chains) in each fraction, when grafted to a surface, was determined from the Δ*D*/-Δ*F* ratio measured by QCM-D at a surface coverage equivalent to -Δ*F* = 2.5 Hz. For the 5^th^ overtone 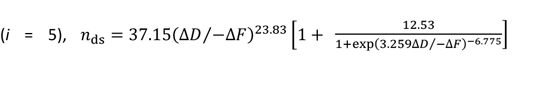 with Δ*D* expressed in units of ppm and Δ*F* in Hz ^50^.

## RESULTS AND DISCUSSION

### Controlling the surface density of anchored molecules through competitive mass-transport limited adsorption - Theory

We consider the interaction scenario depicted in Figure 1: functional molecules each bearing an anchor tag (index 1) are in competition with free anchors (index 2) for specific binding (*i*.*e*., *via* the anchor) to a planar surface. The attachment of each molecule *via* the anchor to the surface is considered strong and irreversible. The main assumption underpinning our approach to control the surface density of the functional molecule is that the rate of binding is limited by the transport of molecules to the surface (mass-transport limited binding) rather than the intrinsic binding rate after their arrival at the surface (kinetically limited binding). Provided that the binders are well mixed, their respective binding rates will depend on their molar concentrations (*c*_1_ and *c*_2_) and rates of diffusion (*D*_1_ and *D*_2_), and also on the conditions of convective fluid transport (if any). Alternative to the rates of diffusion, one may consider the hydrodynamic radii (*R*_1_ and *R*_2_) since *D*1*⁄D*_2_ = *R*_2_*⁄R*_1_ according to the Stokes-Einstein relation. Indeed, any difference in mass between the binders will impact the competitive binding through relative differences in their hydrodynamic radii. Here, we have restricted ourselves to the case of one type of functional molecule mixed with free anchors, but extension to the functionalization of the surface with several distinct molecules is straightforward (*vide infra*).

**Figure 1.**
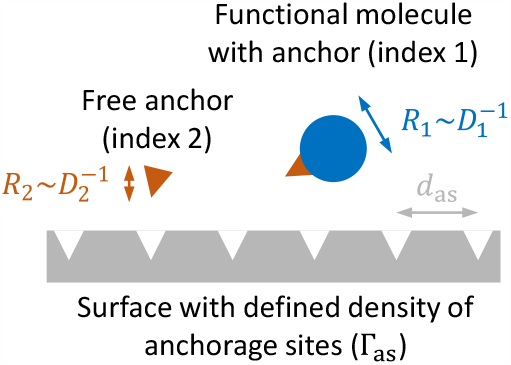
Schematic of the interaction scenario considered in the theory. Main assumptions are that binding is mass-transport limited and irreversible, and that steric hindrance does not impede binding.

### Binding from stagnant solution

A common binding scenario is adsorption from a stagnant solution. The temporal variation in the molar surface density for mass-transport limited binding from a semi-infinite stagnant solution to a planar surface is described by ^37^

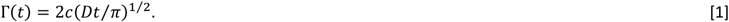

The ratio of the molar surface densities

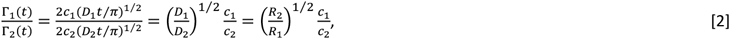

then depends exclusively on the ratio of the concentrations and diffusion constants, and is independent of the incubation time.

We assume additionally that the binding sites on the surface are spaced apart sufficiently for steric hindrance between binders to be negligible. This implies that binding remains mass-transport limited until all binding sites on the surface are occupied. At saturation, the molar surface density of the functional molecule (Γ_1,sat_) and the total molar density of anchorage sites on the surface (Γ_as_) relate as

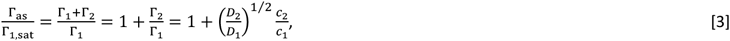

which gives

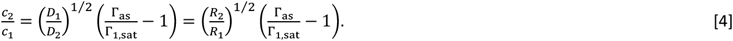

Equation [4] is our main theoretical result. It implies that, for a given Γ_as_, the surface density of the functional molecule can be controlled simply by the molar mixing ratio of the free anchor and the functional molecule. Our approach requires knowledge of the ratio *D*1*⁄D*2, which may be obtained by independent determination of *D*_1_ and *D*_2_, or from the hydrodynamic radii *R*_1_ and *R*_2_ (see Table S1 for an analysis of *D* and *R* for molecules with biotin anchors used in this study). Since the dependence on *D*_1_*⁄D*_2_ (or *R*_2_*⁄R*_1_) is rather weak (exponent 1/2), even a relatively crude estimate should provide satisfactory results.

### Binding with convective fluid transport

Another common binding scenario is adsorption under constant laminar flow. This may be accomplished, for example, through flow in a slit with the target surface being one of the slit walls, or (across a limited surface area) through stirring of the solution in front of the target surface. Irrespective of the exact mechanism of fluid convection, the steady-state rate of mass-transport limited binding is described by ^37^

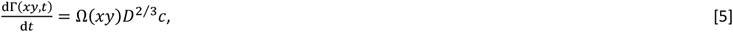

where the parameter Ω(*xy*) encompasses the influence of convective fluid transport, which may depend on the location *xy* on the surface but does not depend on time. If the flow or stirring are sufficiently fast and the concentration of binders in the bulk solution remains essentially unchanged, then steady-state is reached quickly and the binding rate is effectively constant (Γ(*xy, t*) = Ω(*xy*)*D*^2*⁄*3^*ct*) throughout the binding process.

Analogous to the derivation of Eq. [4], one can show that

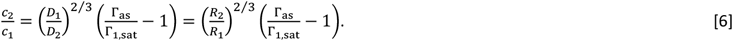

This equation is independent of the exact conditions of convective fluid transport and the surface location, which makes the binding process simple and robust to control.

Comparison of Eqs. [4] and [6] shows that the sensitivity to *D*_1_*⁄D*_2_ (or *R*_2_*⁄R*_1_) is somewhat enhanced yet remains rather weak (power of 2/3) under convective transport. Both equations become identical if the two binders are of similar size (*R*_2_*⁄R*_1_ = *D*_1_*⁄D*_2_ ≈ 1).

### Guidelines for experimental design

The above-described approach is attractive by its simplicity, but makes simplifying assumptions. Rewardingly, these can be met quite easily for a large range of binder sizes (and hence diffusion coefficients) and intrinsic binding rates (provided that binding is effectively irreversible), as defined by the following simple criteria.

First, to satisfy the assumption of mass-transport limited binding implying that the surface acts as a ‘perfect sink’, the flux of binders to the surface should be small compared to the binding rate. This is the case if

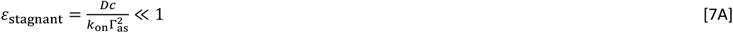

for binding from stagnant solution, and

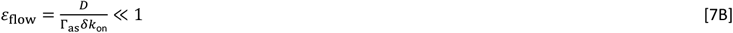

for binding under flow, as described in detail in the Supporting Methods (Sections 1-2) and evaluated in Figures S2-3. Here *k*_on_ is the intrinsic binding rate constant, and *δ* is the thickness of the depletion layer across which molecules from the bulk solution need to diffuse to reach the surface. Relations [7A-B] are useful for experimental design. One can see that *ε*_stagnant_ can be tuned to remain small by limiting the binder concentration, whereas *ε*_flow_ can be tuned to remain small by limiting the flow rate *Q ∝ δ*−^3^. It should here be noted that the liquid layer above the functionalized surface needs to be thicker than the depletion layers for any of the binding molecules, as otherwise the bulk solution is effectively depleted rendering our approach invalid (see the Supporting Methods (Section 1) for details on the determination of *δ*).

Second, if the functional molecules are large (*R*_1_ > *d*_as_), steric hindrance will prevent grafting at densities higher than a maximal surface density that may be significantly smaller than the anchor site density Γ_as_. In such a situation, control of the surface density according to Eqs. [4] and [6] can still be achieved, as long as (i) the diffusion to the surface is unaffected and (ii) care is taken to aim for grafting densities below the maximal value. In practice, 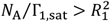 (with *N*A being Avogadro’s number) should provide a reasonable condition for Eqs. [4] and [6] to remain valid.

### Experimental validation of the theoretical predictions – Co-adsorption of two small biotinylated species of similar size

To validate the theoretical predictions, we examined the co-adsorption of two biotinylated molecules of comparable size from stagnant solution. Biotin (*R*_biotin_ = 0.37 nm; Table S1) served as the free anchor, and the biotinylated FITC fluorophore (b-FITC; *R*b_−FITC_ = 0.63 nm; Table S1) as a model target molecule. The receiving surface was a glass-supported lipid bilayer containing biotinylated lipids coated with a dense monolayer of streptavidin (Figure 2A), which we confirmed was homogeneous (Figure S4). The surface was incubated with a mix of free biotin and b-FITC in desired ratios (at constant total binder concentration), and the fluorescence intensity of surface-bound b-FITC at saturation was quantified by confocal microscopy. Using appropriate precautions, the fluorescence intensity is expected to be proportional to, and thus serves as a measure for, the b-FITC surface coverage (see Figure S1).

**Figure 2.**
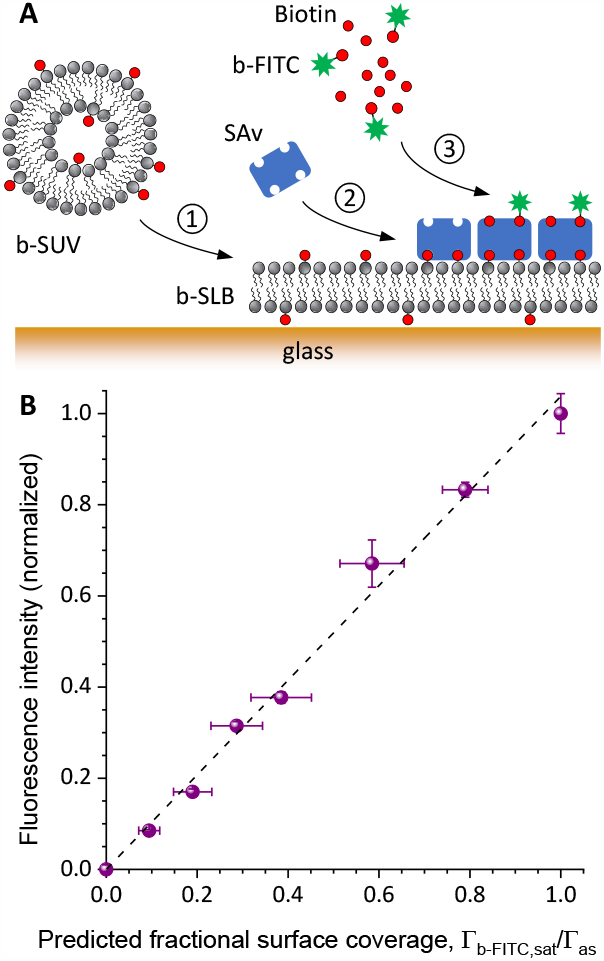
Tuning of the surface densities of two biotinylated species of similar size. **A.** Schematic drawing of the surface functionalisation: *①* SLB formation, *②* streptavidin (SAv) monolayer formation, *③* co-adsorption of b-FITC and biotin. **B**. Fluorescence intensity at saturation, normalized against the intensity at full b-FITC coverage, measured with confocal microscopy for surfaces functionalized with different mixing ratios of b-FITC and biotin. The horizontal axis shows the fractional surface coverage, Γ_b−FITC,sat_*⁄*Γ_as_, of the b-FITC fluorophore (co-adsorbed with free biotin) predicted according to Eq. [4]. The linear trend as demonstrated by the black dashed line through the origin confirms the validity of Eq. [4]. Conditions: c_b−FITC_ + cbiotin = 14 μM was maintained constant, with cb_−FITC_ and cbiotin determined according to Eq. [4] (*ε*_stagnant_ < 0.25).

Figure 2B shows the measured normalized fluorescence intensity as a function of the b-FITC surface density, Γ_b−FITC,sat_, predicted according to Eq. [4] and the molar mixing ratios of b-FITC and biotin. The fluorescence signal obtained with pure biotin was subtracted to eliminate background, and after this correction the fluorescence signal with pure b-FITC was used to normalize all data. The experimental data exhibit a clear linear dependence over the full range of possible surface densities. This demonstrates the validity of Eq. [4].

### Application example 1 – Co-adsorption of two biotinylated species of different size

As a next step, we demonstrate that the method for tuning the surface density also works well for two co-adsorbing molecules with a larger difference in size. We used a biotinylated tandem-repeat of the Z domain of protein A (b-ZZ) as the functional molecule, and biotin as the competing free anchor. The Z domain specifically and stably binds the Fc region of immunoglobulin molecules, making the b-ZZ construct an attractive tool for immunoglobulin isolation and display on surfaces ^51^. It has a hydrodynamic radius substantially larger than biotin (*R*b_−ZZ_ = 2.0 nm; Table S1), implying that the ratio of diffusion constants in Eqs. [4] and [6] is much larger than unity (*R*_b−ZZ_*⁄R*_biotin_ ≈ 5). At the same time, b-ZZ remains small enough to allow occupation of all the available biotin-anchorage sites on a dense streptavidin monolayer (*R*_b−ZZ_ <*d*_as_; Figure 1) ^27^, and it can therefore be expected that steric hindrance does not limit the access of the two co-adsorbing species to the surface. We tested binding from stagnant solution and binding under flow.

### Binding from stagnant solution

For this co-adsorption scenario, spectroscopic ellipsometry (SE) was used to monitor the b-ZZ binding process (Figure 3A). In contrast to fluorescence intensity, SE has the benefit of providing absolute quantification of the surface density of molecules of sufficient size without the need for labels. Streptavidin coated supported lipid bilayers again served as the anchor surface (see Figure S5 for their characterisation by SE). The cuvette-based SE setup was operated in essentially stagnant solution to monitor the binding process (see Methods for details). b-ZZ and biotin were incubated at molar ratios C_biotin_*⁄c*_b-ZZ_ (again at constant total binder concentration) required to obtain the desired fractional b-ZZ surface coverage Γ_b−ZZ,sat_*⁄*Γ_as_ as defined by Eq. [4]. As can be seen in Figure 3B, binding saturated within 10 minutes, and subsequent rinsing in working buffer did not noticeably affect the signal, demonstrating that all b-ZZ was anchored stably.

**Figure 3.**
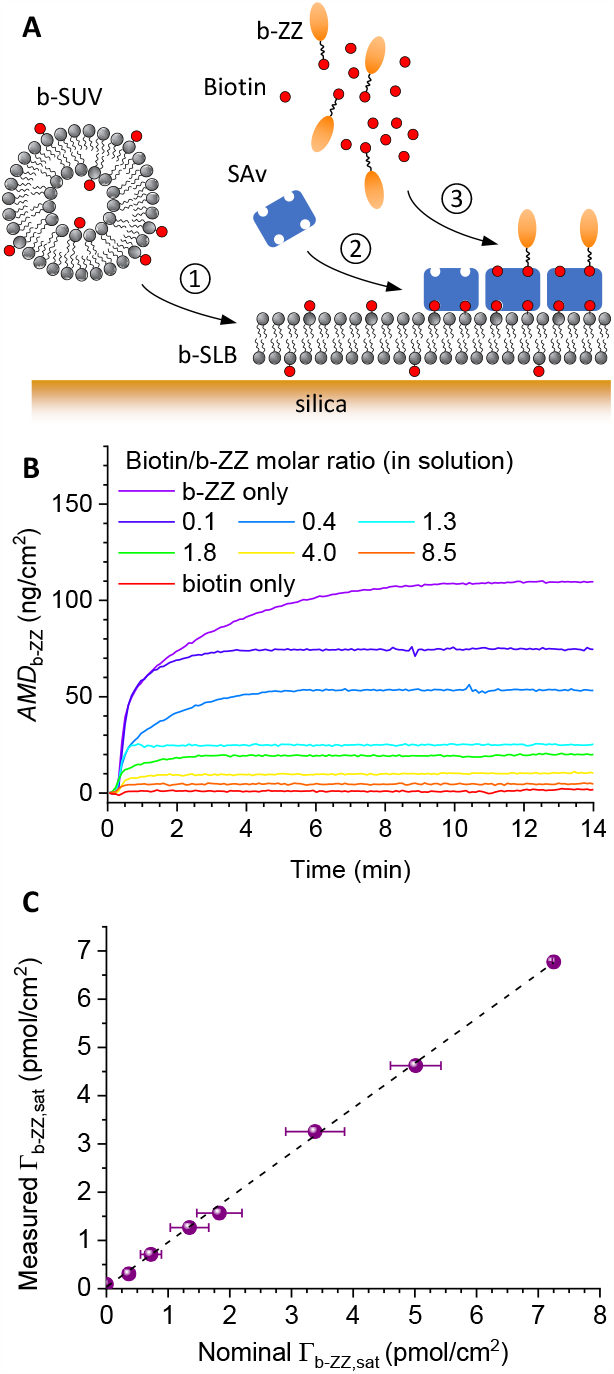
Quantitative tuning of the surface density of two biotinylated proteins of different size (in stagnant solution). **A.** Schematic drawing of the surface functionalisation: ① SLB formation, ② SAv monolayer formation, ③ co-adsorption of b-ZZ and biotin. **B**. Areal mass density of b-ZZ, *AMD*_b-ZZ_, over time, determined by spectroscopic ellipsometry for a range of *c*_biotin_*⁄c*_b-ZZ_ molar ratios (as indicated). Conditions: c_b−ZZ_ + cbiotin = 0.625 μM was maintained constant, with c_b−ZZ_ and cbiotin determined according to Eq. [4] (*ε*_stagnant_ < 0.01); biotin/b-ZZ incubation – 10 min, starting from 0.3 min; see Figure S5 for details of steps① and ②. **C**. b-ZZ surface density at saturation, Γ_b−ZZ,sat_, measured by SE as a function of the nominal b-ZZ surface density predicted according to Eq. [4] from the *c*_biotin_*⁄c*_b-ZZ_ molar ratios and assuming Γ_as_ = 2Γ_SAv_. Error bars along the horizontal axis represent the uncertainty in the concentrations of biotin and b-ZZ when computing Γ_b−ZZ,sat_*⁄*Γ_as_, and the resolution in Γ_SAv_. Error bars along the vertical axis (about the size of the symbols) represent temporal noise and confidence intervals when fitting the SE data. The black dashed line is a linear fit through the data, with a slope of 0.93 ± 0.02.

The b-ZZ surface densities calculated from the SE data at saturation as a function of the nominal b-ZZ surface density are shown in Figure 3C. The nominal surface density was here calculated according to Eq. [4], assuming (somewhat simplistically, *vide infra*) a surface density of anchor sites Γ_as_ = 2Γ_SAv_ . Also, we neglected for simplicity the contribution of the free biotin to the SE signal: the molecular mass of b-ZZ (16.2 kDa) exceeds the molecular mass of biotin (244.3 Da) by far, implying that free biotin makes only a small contribution. The clear linear dependence indicates that the theoretical predictions are indeed consistent with the experimental results.

A linear fit to the data in Figure 3C revealed a slope (0.93 ± 0.02) inferior to one, indicating that the average number of biotin binding sites per streptavidin is 1.86 ± 0.04 rather than 2 (Γ_as_ = 2Γ_SAv_) as one might expect based on the naïve assumption of a symmetric display of streptavidin on the SLB. This stoichiometry is consistent with our previous report (1.74 ± 0.22 for dense SAv monolayers on SLBs) ^27^, which demonstrated that each SAv molecule may anchor *via* either 2 or 3 of its 4 biotin binding sites to the SLB, leading to an average ‘residual valency’ between 2 and 1.

### Binding under flow

We deployed quartz crystal microbalance with dissipation monitoring (QCM-D) to follow the same binding process under flow. The QCM-D flow modules facilitated constant flow over time of reagent solution across the sensor surface. A further benefit of QCM-D was that four experiments could be performed in parallel, thus increasing the data acquisition throughput compared to SE. We confirmed proper formation of the SAv monolayer on SLBs (Figure S6A), and then incubated b-ZZ on its own or mixed with free biotin as the competing anchor. Binding of biotin alone is not detectable by QCM-D, owing to the small size of biotin and its complete burial in the SAv binding pocket ^49^, and the QCM-D responses shown in Figure 4A thus represent exclusively b-ZZ binding. All binding curves show a similar initial binding phase, with a decrease in frequency (Δ*F*; Figure 4A, bottom) demonstrating binding. That the binding rates are approximately constant from shortly after the onset of binding and almost up to saturation, and roughly scale with the concentrations of b-ZZ, is fully consistent with the predictions for steady-state mass-transport limited binding under flow. The concomitant increase in dissipation (Δ*D*; Figure 4A, top) reveals the b-ZZ film to be soft, most likely owing to the flexible peptide linker that connects the biotin anchor to the globular ZZ domain. For b-ZZ alone, the frequency shift at saturation was -23 Hz (Figure S6B), indicating an added film thickness of *h*_b−ZZ_ ≈ 4 nm (see Methods for details) consistent with the estimated hydrodynamic radius of b-ZZ (*h*_b−ZZ_ ≈ 2*R*_b−ZZ_).

**Figure 4.**
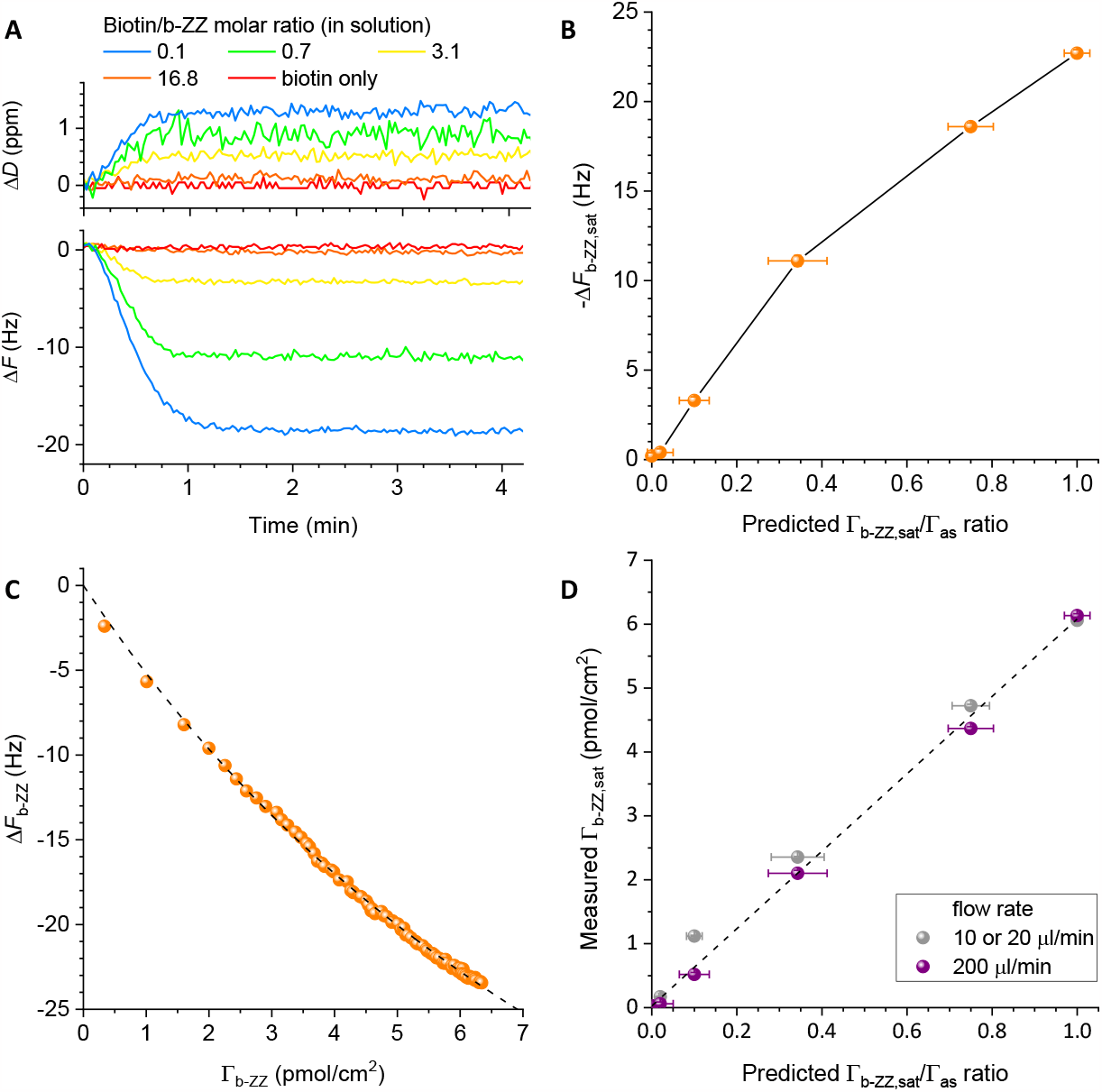
Quantitative tuning of the surface density of two biotinylated proteins of different size (under flow). **A.** QCM-D dissipation shifts Δ*D* (top) and frequency shifts Δ*F* (bottom; for overtone *i*= 5) obtained for b-ZZ mixed with free biotin at distinct molar ratios (as indicated). Conditions: c_b−ZZ_ + cbiotin = 0.625 μM was maintained constant, with c_b−ZZ_ and cbiotin determined according to Eq. [6] (*ε*_flow_ < 0.007); b-ZZ/biotin incubation – 3 min, starting from 0 min; during remaining times, plain working buffer was flown over the sensor surface, flow rate – 200 μl/min. b-ZZ only was incubated at a different flow rate (20 μl/min) and therefore is not displayed in the graph; see Figure S6A for details of the SAv-on-SLB sensor functionalisation and b-ZZ grafting. **B**. Frequency shifts at saturation (Δ*F*_b-ZZ,sat_; *i* = 5) *vs*. the predicted fractional b-ZZ surface coverage (Γ_b−ZZ,sat_*⁄*Γ_as_). **C**. Standard curve relating frequency shifts to b-ZZ molar surface density, obtained through a combined QCM-D/SE experiment (see Figure S7 for details). The dashed line is an empirical fit to the data, with Γ_b−ZZ_ = (*α* − 1)*⁄*[*M*_b−ZZ_(β + *C*^−1^Δ*F*^−1^)] and *α* = 0.8389 ± 0.0012, β = 7.83 ± 0.14 × 10^−4^ cm^2^*⁄*ng, the mass sensitivity constant *C* = 18.0 ng*⁄*(cm^2^Hz), and the molecular mass *M*_b−ZZ_ = 16.2 kDa. **D**. Plot of the measured b-ZZ surface density as a function of Γ_b−ZZ,sat_*⁄*Γ_as_ for two distinct flow rate regimes (200 μl/min - purple spheres; 10 or 20 μl/min – grey spheres, see Figure S6B for details). The black dashed line is a linear fit through the purple data (200 μl/min), with a slope of 6.1 ± 0.2 pmol/cm^2^. b-ZZ surface densities were determined from panel B and Figure S6C, respectively, using the empirical fit from panel C.

The gradual decrease in the magnitude of the frequency shift at saturation, Δ*F*_b-ZZ,sat_, with increasing biotin concentration demonstrates the desired tuning of the b-ZZ surface density (Figure 4B). However, the Δ*F*_b-ZZ,sat_ values cannot be directly translated into surface concentrations because the frequency shift measured by QCM-D not only represents surface bound b-ZZ but also solvent that is hydrodynamically coupled to the protein upon the mechanical shear oscillation of the QCM-D sensor. We and others have previously shown that, for monolayers of globular proteins, the contribution of coupled solvent gradually decreases with protein coverage ^46^-^47, 52^, resulting in a non-trivial dependence of Δ*F*_b−ZZ,sat_ on Γ_b−ZZ,sat_ . We therefore established a standard curve (Figure 4C) to translate the QCM-D frequency shifts into molar surface densities, through an experiment that combined SE and QCM-D analyses *in situ* on the same SAv-on-SLB surface (Figure S7) ^47^. Combining this standard curve with Δ*F*_b-ZZ,sat_ values, we plot in Figure 4D (purple spheres) the inferred Γ_b−ZZ,sat_as a function of the predicted Γ_b−ZZ,sat_*⁄*Γ_as_ ratio. The data demonstrate successful quantitative tuning of the b-ZZ surface density under flow with a linear dependence on the predicted coverage. That the maximal b-ZZ surface density Γ_b−ZZ,max_ = Γ_as_ in the flow-based assay (6.1 ± 0.2 pmol/cm^2^; corresponding to the slope of the linear fit in Figure 4D) was slightly inferior to the density measured in stagnant solution (6.8 ± 0.1 pmol/cm^2^; Figure 3) is likely due to differences in the incubation times of SAv (15 min *vs*. 60 min) affecting the SAv surface density.

We also trialled the same approach in a lower flow rate regime (10 to 20 μl/min instead of 200 μl/min). This also produced a reasonable linear trend (Figure 4D, grey spheres), but with some moderate deviations at a target coverage of 10%. Analysis of the mass transport conditions revealed that the thickness of the depletion layer for the fast-diffusing free biotin approaches the fluid thickness above the QCM-D sensing area in the lower flow rate regime (see Supporting Methods (Section 1)). A likely explanation for the larger-than-expected b-ZZ binding at intermediate target coverages therefore is excessive depletion (and thus reduced competition) of free biotin from the bulk solution. This example demonstrates the importance of an appropriate design of incubation conditions to achieve the desired surface densities.

### Application example 2 – Tuning the surface density of the receptor P-selectin through an adapter protein

The previous example demonstrated direct control over the surface density and presentation of a functional molecule *via* a single, site-specific biotin anchor. A large variety of methods for biotinylation exist making this approach potentially useful for many functional molecules of interest. However, it is sometimes impractical or technically challenging to equip the molecule of interest with a single biotin anchor at a specific site. In these instances, more complex strategies for surface anchorage are required. Here, we demonstrate how the surface density of a recombinant receptor protein can be controlled with the help of an adapter protein.

P-selectin (CD62P) is a transmembrane receptor expressed at the surface of activated endothelial cells lining blood vessels. P-selectin mediates the adhesion and rolling of leukocytes at the blood vessel wall through interaction with its ligand PSGL-1 ^53^, which is an important step of the migration of circulating immune cells into interstitial tissue. The display of ectodomains of cell adhesion receptors on artificial surfaces is an attractive route for biophysical analyses of the molecular and physical mechanisms of cell adhesion. In this context, the ability to anchor receptors at defined surface densities is particularly pertinent. The density of P-selectin receptors on endothelial cells, for example, has been estimated to about 350 per μm2, corresponding to a root mean square distance between receptors of approximately 50 nm ^54^. The density of biotin-binding sites on our densely packed streptavidin monolayer, on the other hand, is approximately 1/(5.0 nm)^2^, or 40,000 per μm^2^. The large difference illustrates the need for quantitative tuning to bring the model system closer to the biological conditions.

Here, we demonstrate the quantitative tuning of the surface density of P-selectin receptors through the surface density of the b-ZZ adapter protein (Figure 5A). The recombinant receptor construct was a fusion of the ectodomain of human P-selectin and the Fc part of human IgG1 immunoglobulin which binds the Z domains of b-ZZ. Owing to the dimeric nature of the Fc part, each P-selectin-Fc molecule contains two P-selectin ectodomains. The surface density of b-ZZ was tuned as described in the previous example, and P-selectin-Fc was then added at the same final concentration (67 nM) irrespective of b-ZZ coverage.

**Figure 5.**
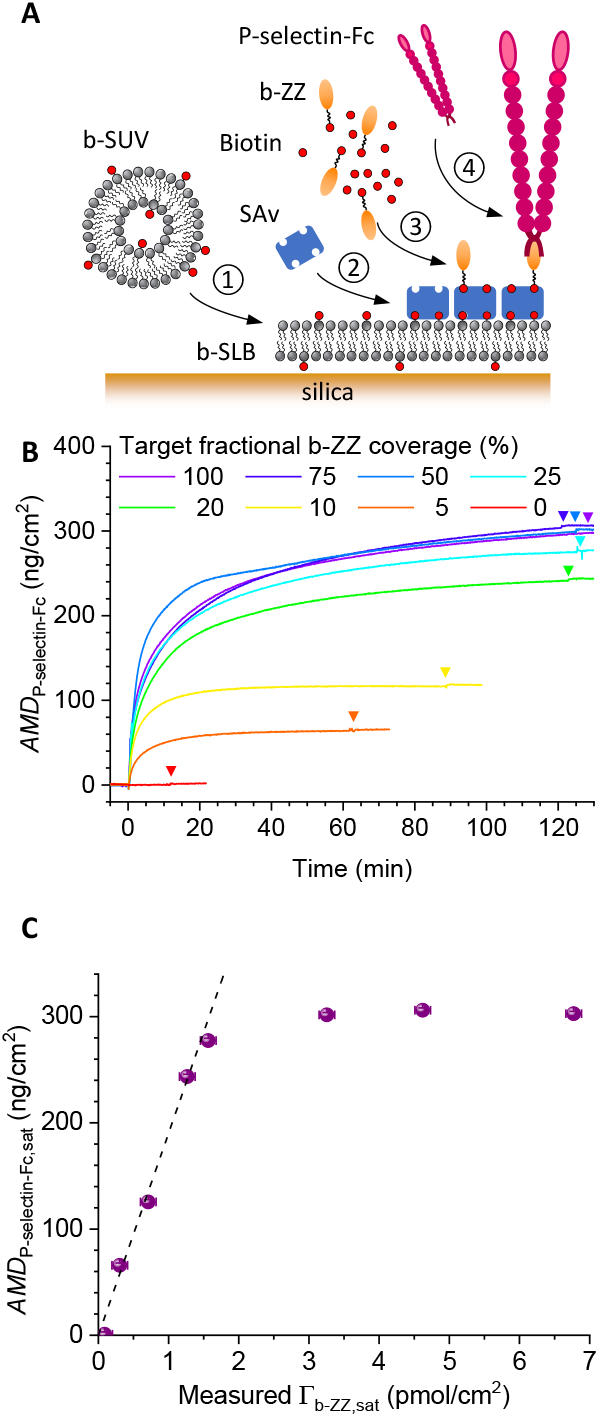
Tuning the surface density of P-selectin receptors through the surface density of b-ZZ adapter protein. **A.** Schematic drawing of the surface functionalisation: ① SLB formation, ② streptavidin monolayer formation, ③ co-adsorption of b-ZZ and biotin, ④ P-selectin-Fc anchorage. **B**. Areal mass density, *AMD*, of P-selectin over time, determined by SE for a range of fractional b-ZZ surface coverages (as indicated). Conditions: P-selectin-Fc (67 nM) was incubated at 0 min; the start of rinses with working buffer is indicated by arrowheads in matching colours. **C**. Areal mass density of P-selectin at saturation or after 2 hours of incubation (derived from B) as a function of the measured b-ZZ surface density (taken from Figure 3C). Error bars along both axes are about the size of the symbols and represent temporal noise and the effects of confidence intervals and parameter correlations when fitting the SE data for b-ZZ and P-selectin-Fc binding, respectively. The black dashed line is a linear fit through the data for Γ_b−ZZ,sat_≤ 1.5 pmol*⁄*cm^2^.

Figure 5B shows the kinetics of P-selectin-Fc binding to surfaces displaying b-ZZ at defined densities. For sufficiently low surface densities of adapter protein (*i*.*e*., less than 20% of maximal coverage), P-selectin binding saturated within less than 1 h of incubation. For higher b-ZZ surface densities, an initial fast binding phase was followed by a second slow binding phase, and binding did not reach full saturation even after 2 h of incubation. Reassuringly, all binding responses remained unchanged upon rinsing in working buffer, and binding to 0% b-ZZ surfaces was essentially absent, demonstrating specific and stable anchorage of P-selectin-Fc to b-ZZ *via* the Fc/Z interaction. The initial fast binding phase illustrates the challenge of controlling the grafting density on the surface through the tuning of the incubation time on a 100% b-ZZ surface.

Figure 5C shows the maximal P-selectin areal mass density as a function of the b-ZZ surface density at saturation. The data for the four lowest b-ZZ coverages (including the control with 0% b-ZZ) show a very good linear dependence, and thus demonstrate how surfaces with a tuneable density of adaptor proteins (here b-ZZ) enable quantitative control over the surface density of the target receptor (here P-selectin).

Another salient feature of Figure 5C is the plateau in P-selectin-Fc coverage at higher b-ZZ surface densities, which we attribute to a dense protein monolayer. That the transition to the plateau (around 25% of maximal b-ZZ coverage, equivalent to 1.6 pmol/cm^2^) coincides with the emergence of a slow P-selectin-Fc binding phase (Figure 5B; indicating steric hindrance) is consistent with this explanation. Moreover, an anchor site surface density of 1.6 pmol/cm^2^ is equivalent to an average surface area of 110 nm^2^ per b-ZZ. This value is reasonable for an effective cross section of P-selectin-Fc, considering that P-selectin-Fc molecules are homodimers, and that P-selectin ectodomains are elongated with typically 40 nm in length and a few nm in diameter ^55^.

### Application example 3: Post-chromatographic analysis of the products of an anchor ligation reaction

In all examples so far, we exploited known mixing ratios of functional molecules and free anchors (biotin) to tune the surface density of the functional molecule. In some cases, it is instead of interest to determine the mixing ratio from the measured surface density of the functional molecule. A case in point is the analysis of contamination of a sample with free anchors following an anchor ligation reaction. We here demonstrate this for the site-specific biotinylation of glycosaminoglycans (GAGs).

GAGs are linear carbohydrate polymers ubiquitous on cell surfaces and in extracellular matrices, and contribute to a wide range of cell and tissue functions, including in tissue development, inflammation and immunity ^56^. Isolated from natural sources, GAG preparations typically have a high size dispersity, and a heterogeneous composition notably with regard to the level of sulfation of the constituent monosaccharides. The chemical modification of a single end of GAGs is often desirable (*e*.*g*., with a biotin that can be anchored to biotin-binding proteins for functional molecular and cellular interaction assays), but the compositional complexity of GAGs make the analysis of the reaction products challenging with conventional methods such as nuclear magnetic resonance or mass spectrometry. We here consider the biotinylation of GAGs at their reducing end by oxime ligation ^49^. As in application example 2, we deploy QCM-D with streptavidin-coated SLBs to analyse the reaction product for their content in biotinylated GAGs (GAG-b) and in residual unreacted biotin anchor contaminants.

### Theoretical considerations

For films of end-grafted GAGs, and in contrast to the globular proteins (such as b-ZZ; Figure 4), the QCM-D frequency shift Δ*F* is proportional to surface coverage Γ to a good approximation49. Moreover, the alkoxyamine-modified biotin used for biotinylation only entails a very small QCM-D response (−Δ*F*_b−alkoxyamine_ ≤ 0.5 Hz; Figure S8A). We can thus take Δ*F*_pure,sat_ to be the response at saturation for a GAG-b film formed from a pure solution of biotinylated GAGs, and Δ*F*_sample,sat_ to be the response at saturation for a GAG-b film formed from a GAG-b solution contaminated with free biotin. In this Case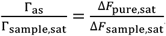, and Eq. [6] becomes

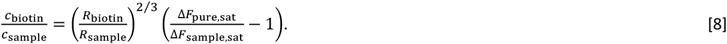

Equation [8] can be used to quantify the molar ratio of free biotin to biotinylated GAGs from the measured frequency shift for a contaminated and a pure sample, with reasonable assumptions about the size of the GAG molecules. This method is particularly attractive for post-chromatographic analysis of sample concentration and anchor contamination, as will be shown in the following.

### Analysis of biotin anchor contamination in a representative GAG sample

A commercial sample of chondroitin sulfate D (CS-D) GAGs was reacted with alkoxyamine-modified biotin for site-specific biotinylation of the GAG’s reducing end ^49^. Figure 6 presents the analysis of the biotinylated CS-D GAGs following size-exclusion chromatography of the reaction products. The QCM-D time traces upon GAG-b incubation (Figures 6B and S8C) featured the saturable binding responses expected for monolayer formation in eluate fractions 2 to 8, demonstrating that these fractions contained biotinylated CS-D. Whilst the binding rates differed between fractions, they typically varied little throughout most of the binding processes up until close to saturation (Figure S8C), consistent with steady-state mass-transport limited binding. Binding from fraction 3 was fastest and also reached the highest coverage at saturation (Figure S8C), indicating that this fraction was purest and the most concentrated. There was no binding in fraction 1 (Figure S8C), indicating that this fraction contained no GAGs.

**Figure 6.**
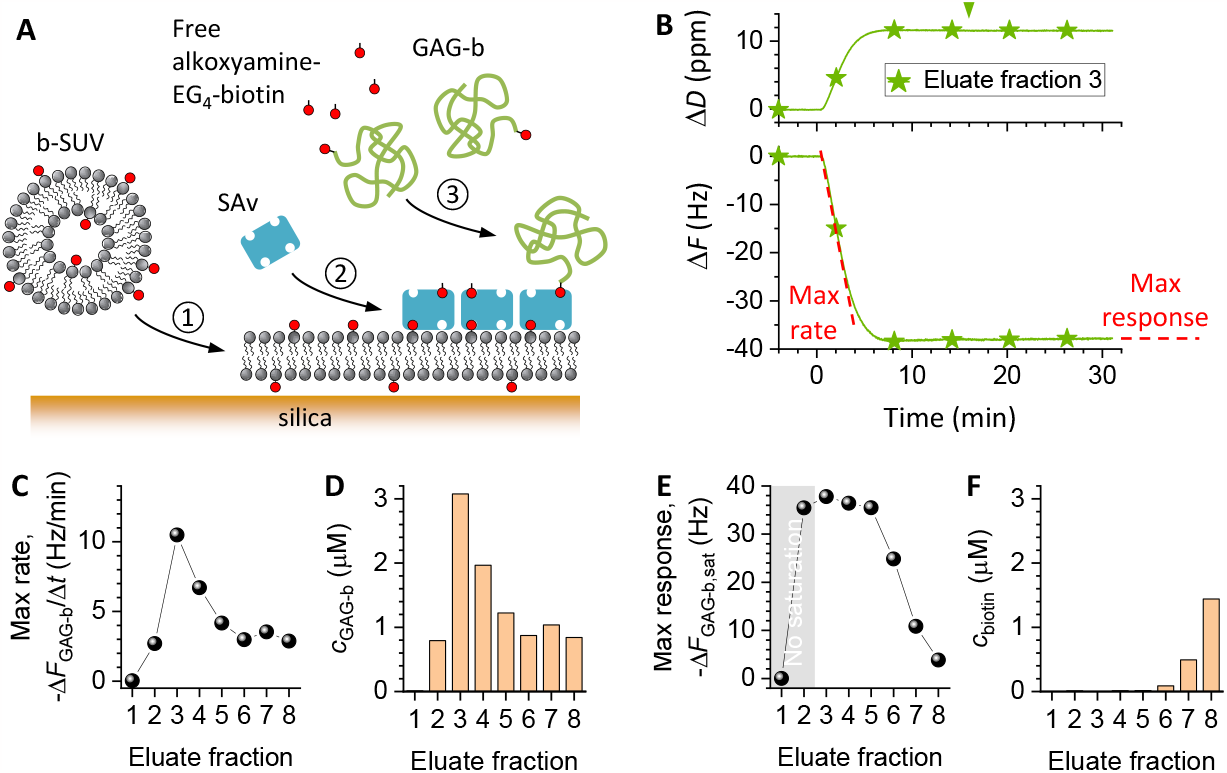
Analysis of concentration and anchor contamination of GAG samples. **A.** Schematic showing the surface functionalisation steps for post-chromatographic QCM-D analysis of the GAG samples: ① SLB formation, ② SAv monolayer formation, ③ binding of biotinylated GAGs and/or free alkoxyamine-EG4-biotin. **B**. QCM-D responses (Δ*F* - bottom; Δ*D* - top; *i* = 5) for the binding of biotinylated CS-D GAG (GAG-b) from eluate fraction 3. Data for all other eluate fractions (EF) are shown in Figure S8C. Conditions: GAG-b incubation - started at 0 min, and proceeded for 16 min (during other times, plain working buffer was flown over the sensor); flow rate – 20 μl/min; GAG-b concentration – the eluate fractions as retrieved from the size-exclusion column were diluted 15-fold for QCM-D analysis. The steepest slope in the Δ*F*GAG_-b_ *vs*. time graph (Max rate, -Δ*F*_GAG-b_/Δ*t*; indicated by the tilted red dashed line) reflects the rate of steady-state mass-transport limited binding. The frequency shift at saturation, Δ*F*_GAG-b,sat_ (Max response; indicated by the horizontal red dashed line), is a measure of the fraction of biotin binding sites occupied by biotinylated GAGs. **C**. Max rate, -Δ*F*_GAG-b_/Δ*t*, as a function of the eluate fraction. **D**. GAG-b concentration in the eluate fractions, calculated from -Δ*F*_GAG-b_/Δ*t* and reference data for another GAG-b of known concentration and similar size (see Figure S8B for details). **E**. Max response, -Δ*F*_GAG-b,sat_, as a function of the eluate fraction. The grey background highlights the fractions for which binding did not saturate during the set incubation time. **F**. Concentration of residual free alkoxyamine-EG4-biotin in the eluate fractions, estimated through Eq. [8] based on the data in D and E, and *R*_GAG−*b*_*⁄Rb*_−alkoxyamine_ ≈ 12.

To quantify the GAG-b concentration profile across the fractions, we extracted the slopes of highest magnitude, -Δ*F*_GAG-b_/Δ*t*, from the frequency shift *vs*. time traces (Figure 6B-C). According to Eq. [5], the rate of binding is proportional to the GAG-b concentration. The slopes therefore directly report on the relative concentration differences, and thus revealed that fractions 3 to 5 contain most GAG-b. Considering that all fractions had the same volume, we can estimate through integration that fractions 3 to 5 contained 64% of the total GAG-b content in the 8 fractions.

Δ*D*/-Δ*F* ratios are very sensitive to the degree of polymerisation of surface-grafted GAGs, and can be used to quantify the mean GAG contour length, as we have recently shown ^50^. Analysis of the Δ*D*/-Δ*F* ratios (Figure S8D) revealed effective GAG sizes in the range of *n*_ds_ = 77 to 90 disaccharides largely independent of the eluate fraction (Figure S8D, inset). Reference data from approximately length-matched GAG-b (hyaluronan with 95 ± 5 disaccharides; Figure S8B) of known concentration enabled us to estimate the GAG-b concentration in each fraction (Figure 6D).

To assess the degree of contamination with free biotin, we analysed the magnitude of the frequency shift at saturation, −Δ*F*_GAG−b,sat_, across the fractions (Figure 6B, E). Across fractions 6 to 8, −Δ*F*_GAG−b,sat_ gradually decreased indicating increasing contamination with free biotin reactant, consistent with the expected late elution of the comparatively small alkoxyamine-EG4-biotin molecules from the size exclusion column. In contrast, −Δ*F*_GAG−b,sat_ was largest and essentially constant across fractions 3 to 5. The occurrence of a plateau here indicates that these fractions are essentially devoid of free biotin. A constant level of free biotin contamination across these fractions is unlikely because alkoxyamine-EG4-biotin is expected to elute as a relatively sharp peak. It is expected that fraction 2 is also pure, even though this fraction did not quite reach saturation within the limited incubation time owing to its low concentration and binding rate (Figure S8C).

Lastly, we used Eq. [8] with the data in Figures 6D-E to estimate the molar concentration of free alkoxyamine-EG4-biotin in each eluate fraction (Figure 6F). To this end, the hydrodynamic radii were estimated at *R*_GAG−b_ ≈ 7. .8 nm (for *n*ds = 77..90) and *R*_b−alkoxyamine_ ≈ 0.6 nm, respectively (Table S1). We note that the values for eluate fractions 6 to 8 are potentially affected by a flow rate being too slow such that the depletion layer thickness exceeds the chamber height for the free alkoxyamine-EG4-biotin. The concentration of free alkoxyamine-EG_4_-biotin may therefore be somewhat underestimated (by up to a factor of 2, based on the results in Figure 4).

Taken together, the post-chromatographic analysis of sample binding to a biotin-capturing surface thus enabled quantitation of the GAG-b concentration and the contamination with free biotin reactant across the eluate fractions.

### Workflow to tune ligand densities through competitive anchorage

We have demonstrated how competitive mass-transport limited anchorage can be exploited to quantitatively tune ligand grafting densities (application examples 1 and 2) and to quantify the contamination of a solution of anchor tagged molecules by free anchor reactants (application example 3). The combination of relatively simple theories, and their validation by experiments and numerical simulations, has led to a set of guidelines. To facilitate the adoption of our method to tune ligand densities through competitive anchorage, we here recapitulate the main elements of the workflow:

1. Determine the concentrations of the anchor-tagged ligand and the free anchor in their stock solutions, as well as the hydrodynamic radii *R*_1|2_ (or diffusion constants *D*_1|2_) of the two molecules. For folded proteins, for example, good approximations can be found from their molecular mass and/or radius of gyration ^57^.
2. Determine the density of anchor sites Γ_as_ on the target surface.
3. Define the desired surface density of the anchor-tagged ligand (Γ_1,sat_).
4. Define the incubation conditions, and calculate the required concentration ratio *c*2*⁄c*1according to Eq. [4] (for stagnant solution) or Eq. [6] (for flow).
5. Tune the incubation conditions such that the relevant *ε* parameter remains sufficiently small. For stagnant solution (Eq. [7A]), this is most easily done through the binder concentration, whereas for flowing solution (Eq. [7B]) it is most easily done through the flow rate.
6. Ascertain that the liquid above the surface remains thicker than the depletion layers, such that excessive depletion of the bulk solution is avoided (Eq. [S4] for stagnant solution, Eq. [S5] for flow in a slit).
7. Mix and incubate the binders as per the conditions defined in steps 4 to 6, and incubate until saturation. The required incubation times can be estimated from Eq. [1] (for stagnant solution) and Eqs. [5] or [S5] (for flow).

By examining different scenarios of the binding process (stagnant solution *vs*. flow, similar *vs*. different sizes of the co-adsorbing molecules) we have demonstrated that the proposed approach can be applied quite broadly. Our approach is relatively simple and robust. It can be readily implemented in a variety of devices including for label-free surface-based interaction analysis (*e*.*g*., by QCM-D, SE, or surface plasmon resonance), and in well plates or microfluidic devices for further microscopic or spectroscopic analyses.

We note in passing that the theoretical estimates required for steps 5 and 6 in the above workflow can be replaced by control experiments in the case of binding under flow. Under the appropriate mass-transport limited conditions, the steady-state binding rate should scale with the volumetric flow rate as dΓ*⁄*d*t ∝ Q*^1⁄3^ ^37^. In contrast, the dependence of the binding rate on flow rate should vanish (dΓ*⁄*d*t ∝ Qv* with *v* = 0) if binding is entirely kinetically limited, or at least decrease (0 < *v* < 1*⁄*3) if intrinsic binding rates and mass-transport jointly limit binding. Conversely, depletion of binders from the bulk solution (*e*.*g*., owing to too thin a slit) should increase the dependence of the binding rate on the flow rate (*v* > 1*⁄*3). Experimental determination of the power *v* thus provides an alternative route to test if binding conditions are appropriate. This can be helpful, for example, in cases where the flow geometry is complex or unknown, or where *k*_on_ is unknown.

To further assess its usefulness, we have analysed the performance errors of our competitive binding method. With reasonable assumptions (see Supporting Methods (Section 3) for details), the relative errors remain in the range of 10% irrespective of the targeted level of ligand surface density (Figure S9). For comparison, kinetically controlled binding and depletion controlled binding methods tend to have higher errors, and particularly so for low ligand surface densities (Figure S9).

### Functionalising surfaces with multiple binders

In the presented examples, we have limited ourselves to controlling the surface density of one type of ligand at a time (FITC, ZZ or P-selectin). However, the approach can be readily extended to two or more ligands with the same anchor tag. This makes the method attractive for the creation of more complex surfaces, for example, to mimic biological interfaces such as the cell surface.

In the most general case, one wishes to graft *N* − 1 distinct ligands to a surface by co-incubation with the free anchor (*N* species in total). The target surface densities Γ_1_ to Γ_*N*−1_ are fixed, and the surface density of the free anchor is given by 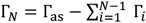. The incubation concentrations *ci* can then be calculated for two different yet practically relevant cases.

If the concentration of one of the binders (here taken as *i* = 1) is fixed, then the concentration of any other binder *i* is given by (see Eq. [2])

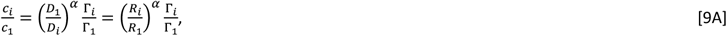

with *α* = 1/2 in stagnant solution, and *α* = 2/3 under flow.

If the total concentration of binders 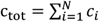 is fixed, then the concentrations of any two binders *i* and *j* relate as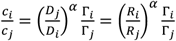. Taken together, these two equations give

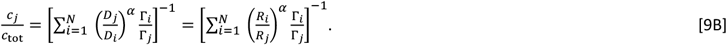

### Expansion to other anchors

For the examples presented here, we deployed biotin as the anchor tag. The method, however, is readily applicable for other anchor tags. The main constraints on the anchor are that its intrinsic binding rate is sufficiently high so as to facilitate mass-transport limited binding and that binding is effectively irreversible. For nickel chelation of polyhistidine tags ^58^, and for IgG binding to protein A or protein G ^59^, binding rates on the order of 105 M^−1^s^−1^ have been reported. The intrinsic binding rate for DNA hybridization can also reach values as high as 105. . 106 M^−1^s^−1^. Although these values are somewhat lower than the intrinsic biotin/streptavidin binding rate (*k*_on_ ≈ 107 M^−1^s^−1 60^) the *ε* values can be kept small according to Eqs. [7A-B] to retain mass-transport limited binding conditions. We should here also note that the attachment of biotin (or any other small anchor tag) to a large binder is expected to decrease *k*on to some extent, owing to the extra time required for rotation of the binder to position its anchor tag appropriately for anchorage to the surface. Fortunately, our method is insensitive to variations in *k*_on_, as long as the binding process remains mass-transport limited.

### Expansion to other anchorage platforms

All application examples provided in this paper relied on the same anchorage platform, a supported lipid bilayer with a dense monolayer of streptavidin. This choice was made out of convenience: the authors have extensive experience with this platform, which is versatile to anchor biotinylated molecules with little or no non-specific binding ^27-28^. The here-presented method to tune binder grafting densities, however, should also be applicable to many other platforms. First, it should be possible to replace streptavidin by other biotin binding proteins with similarly high intrinsic binding rates, such as neutravidin or traptavidin ^61^. Second, whilst a fluid supported lipid bilayer enabled in-plane diffusion of streptavidin (Figure S4) in our platform, such anchor site mobility is not required for our method to work. For example, the method should also work for biotin-binding proteins attached to biotinylated SAMs ^27-28^ or directly coupled to a surface ^62^. Third, the method should work just as well for surfaces with a reduced surface density of anchorage sites (Γ_as_), even if a high Γ_as_ maximises the range of accessible binder surface densities and the range of suitable experimental conditions (*cf*. Eqs. [7A-B]). A reduced surface density of anchorage sites would be beneficial, for example, to ensure full lateral mobility of binders on fluid supported lipid bilayers wherever this is required for the target application (a high streptavidin surface density can entail two-dimensional crystallisation of streptavidin, effectively impairing lateral mobility ^63^).

### Building in added passivation

In our examples, we have considered free anchors as competitors. However, other competitors may be considered as well and can serve to further enhance the functionality of the surface. For example, instead of free biotin one may use biotin functionalized with an inert polymer such as oligoethylene glycol to further enhance the non-fouling properties of the surface ^20^. When choosing the passivating molecule, one should ensure that it is small enough to occupy all binding sites on the surface so that steric hindrance does not skew the expected grafting density.

## CONCLUSIONS

We have established a robust and versatile method to control the grafting density of ligands on a platform presenting binding sites for an anchor such as biotin. The theory developed permits to easily calculate the mixing ratio between anchor-tagged ligands and free anchors that will provide the desired ligand surface density as a function of the incubation conditions. We have provided guidelines to ensure that the conditions for accurate control of the surface density are met, and experimental demonstration of this approach in model cases of surface functionalization and purity control of complex molecules. Our method opens new avenues to develop biomimetic model surfaces where grafting of one or more complex molecules to a surface at controlled densities is required.

## SUPPORTING INFORMATION

Supporting theoretical methods, supporting table of salient binder properties (Table S1), supporting experimental and theoretical figures (Figures S1 to S9) and associated supporting references.

## ACKNOWLEDGEMENTS

This work was financially supported by the BBSRC (Research Grant BB/T001631/1, Research Grant BB/X00158X/1, and Partnering Award BB/W018500/1; to R.P.R.) and the Leverhulme Trust (RPG-2018-100, to J.C.F.K. and R.P.R.). O.K. acknowledges financial support from LabEx Tec21 (Investissements d’Avenir, ANR-11-LABX-0030), and for access to the confocal microscope facility. D.D. and L.B. are members of the LabEx Tec21. L.B. acknowledges financial support from the Centre National d’Etudes Spatiales (CNES) through the Endothelial Dysfunction (DYSFENDOT) program.

## AUTHOR CONTRIBUTIONS

O.K., S.S. and X.Z. designed experiments, collected and analysed data, and drafted the original manuscript. A.R.E.R. collected data. L.C.-G. designed experiments. J.C.F.K. designed experiments, supervised the project and raised funds. L.B. and D.D. conceived the study, designed experiments, analysed data, performed numerical analyses, drafted the original manuscript, supervised the project, and raised funds. R.P.R. conceived the study, designed experiments, analysed data, drafted the original manuscript, supervised the project, and raised funds. All authors contributed to manuscript review and editing.

## ToC graphic

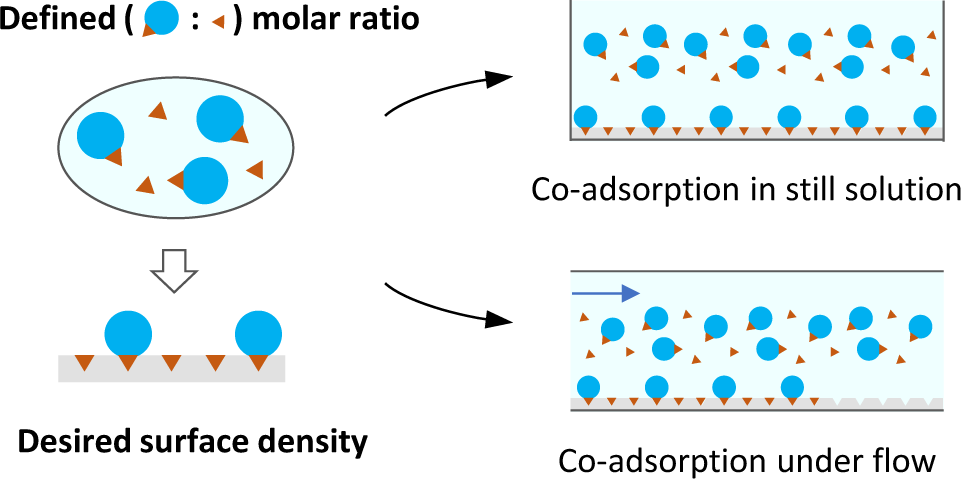

## Notes

### Competing Interest Statement

The authors have declared no competing interest.

### Summary of Updates

Minor revisions of the main text, and expansion of the discussion.

