## Supporting Information for "Competitive specific anchorage of molecules onto surfaces: quantitative control of grafting densities and contamination by free anchors"

**Table of contents**

Supporting Methods S2

Supporting Table S8

Supporting Figures S9

Supporting References S17

### SUPPORTING METHODS

#### 1. Analysis of the validity range of Eqs. [4] and [6]

This section provides a detailed analysis of the validity range of the 'perfect sink' approximation underpinning Eqs. [4] and [6].

***Mass-transport limited binding.*** The assumption of mass-transport limited binding implies that the surface acts as a perfect sink. That is, all molecules that arrive at the surface rapidly bind and the concentration near the surface remains zero at all times throughout the binding process. This approximation obviously fails as the surface density of anchors approaches saturation. In this situation, the solute concentration near the surface will rise over time and ultimately approach the bulk concentration. Moreover, the assumption of mass transport limited binding may fail well below saturation, if the intrinsic binding rate $k_{\text{on}}$ is relatively low.

Equations [4] and [6] are strictly correct for any binder size and intrinsic binding rate, if the two binders have the same hydrodynamic size ($R_2/R_1 = D_1/D_2 = 1$) and the same intrinsic binding rate ($k_{\text{on},1}/k_{\text{on},2} = 1$), irrespective of whether the perfect sink approximation is met throughout the adsorption process. In fact, under these particular conditions, mass-transport limited binding and kinetically limited binding yield the same relation, $c_2/c_1 = \Gamma_{2,\text{sat}}/\Gamma_{1,\text{sat}}$. Such stringent conditions, however, are difficult to meet in experiments and this simple relation thus has limited practical use.

If the binders differ in size, then the validity of the perfect sink approximation can be assessed by comparing the time scale of binding

$$\tau_{\text{binding}} = \frac{1}{k_{\text{on}} c} \qquad \text{[S1]}$$

with the time required to transport enough molecules to the surface to saturate all the anchorage sites. The latter differs for stagnant solutions and convective fluid transport. In stagnant solution, the transport time is obtained from Eq. [1] as

$$\tau_{\text{transport,stagnant}} = \frac{\pi}{4}\frac{\Gamma_{\text{as}}^2}{Dc^2} \sim \frac{\Gamma_{\text{as}}^2}{Dc^2}, \qquad \text{[S2A]}$$

where we drop the numerical pre-factor $\pi/4$ for simplicity. Convective fluid transport leads to the formation of a surface-proximal layer that is depleted in binders. At quasi-steady state, the concentration profile and the depletion layer thickness $\delta$ are constant in time (but may depend on the location $xy$ on the surface). Binders need to diffuse through the depletion layer to reach the surface, and the binder surface density is described by $\Gamma(xy,t) \approx \frac{D}{\delta(xy)} ct$. This yields the transport time

$$\tau_{\text{transport,flow}} = \frac{\Gamma_{\text{as}}\delta(xy)}{Dc}. \qquad \text{[S2B]}$$

Equations [S1] and [S2A-B] lead to the ratios

$$\varepsilon_{\text{stagnant}} = \frac{\tau_{\text{binding}}}{\tau_{\text{transport,stagnant}}} = \frac{4}{\pi}\frac{Dc}{k_{\text{on}}\Gamma_{\text{as}}^2} \sim \frac{Dc}{k_{\text{on}}\Gamma_{\text{as}}^2} \qquad \text{[S3A]}$$

and

$$\varepsilon_{\text{flow}} = \frac{\tau_{\text{binding}}}{\tau_{\text{transport,flow}}} = \frac{D}{\Gamma_{\text{as}}\delta k_{on}}, \qquad \text{[S3B]}$$

which were given in Eqs. [7A-B]. The perfect sink approximation is equivalent to $\varepsilon \to 0$. How well Eqs. [4] and [6] approximate real experimental conditions can be analyzed through numerical simulations, as described further below. These analyses demonstrate that:

- Equation [4] is valid with a relative error in $\Gamma_{1,\text{sat}}$ inferior to 10% provided that $\varepsilon_{\text{stagnant}} < 0.1$, and that the ratio $D_1/D_2$ (or $R_2/R_1$) lies between 0.1 and 10 (Figure S2E). Decreasing $\varepsilon_{\text{stagnant}}$ further ensures better accuracy in the grafting density control, even for large differences in hydrodynamic radii (Figure S2C-D).

- Equation [6] is valid with a relative error in $\Gamma_{1,\text{sat}}$ inferior to 15% provided that $\varepsilon_{\text{flow}} < 0.01$, and that the ratio $D_1/D_2$ (or $R_2/R_1$) lies between 0.1 and 10 (Figure S3D).

We have also tested that the predicted values for the surface density of binder 1 are robust against a possible pitfall. Grafting of the anchor moiety to the functional molecule may reduce the intrinsic binding rate $k_{\text{on},1}$ compared to the free anchor ($k_{\text{on},2}$), for example, by restricting the rate of thermal rotation. Provided that $\varepsilon$ remains sufficiently small for both binders, this difference in binding rate does not appreciably influence the final surface density of binder 1 (Figures S2F-H and S3C-E).

Conditions for mass-transport limited binding are readily met for high affinity interactions, such as the interaction of biotin with streptavidin, as shall be illustrated through two worked examples.

***Worked example for binding from stagnant solution.*** We consider interactions between biotin and streptavidin, for which $k_{\text{on}} = 10^7\ \text{M}^{-1}\text{s}^{-1}$ [1]. Considering that any biotinylated molecule will diffuse at a rate that is smaller than plain biotin ($D \leq 580\ \mu\text{m}^2/\text{s}$), and taking $\Gamma_{\text{as}}$ = 6.8 pmol/cm$^2$ (Figure 3C), we estimate $\varepsilon_{\text{stagnant}} < 0.02\frac{c}{\mu\text{M}}$. This analysis shows that mass-transport limited competitive binding is readily achieved with sub-µM concentrations of biotinylated binders.

The thickness of the depletion layer in stagnant solution for each binder $i$ increases with time as $\delta_{i,\text{stagnant}} \sim \sqrt{D_i t}$. We can use the surface saturation time $\tau_{\text{stagnant,max}}$ to estimate the maximal extension $\delta_{i,\text{stagnant,max}}$ of this layer. At this time, the surface density of binder $i$ is given (Eq. [1]) by $\Gamma_i = 2c_i\sqrt{\frac{D_i \tau_{\text{stagnant,max}}}{\pi}}$, with $\sum_{i=1}^{N} \Gamma_i = \Gamma_{\text{as}}$. Combining these equations gives $\sqrt{\tau_{\text{stagnant,max}}} = \frac{\Gamma_{\text{as}}}{\sum_{i=1}^{N} 2c_i\sqrt{\frac{D_i}{\pi}}}$ and, eventually,

$$\delta_{i,\text{stagnant,max}} \sim \frac{\sqrt{\pi}}{2}\frac{\Gamma_{\text{as}}\sqrt{D_i}}{\sum_{j=1}^{N} c_j\sqrt{D_j}}, \qquad \text{[S4A]}$$

To match the assumption of an infinite reservoir of molecules in solution, one should ensure that $\delta_{i,\text{stagnant,max}} < H$, with $H$ the thickness of the liquid reservoir in the direction perpendicular to the surface being functionalized. A simpler (albeit less stringent) expression is obtained when considering the upper bound of the numerator and the lower bound of the dominator in Eq. [S4A], as

$$\delta_{\text{stagnant,max}} \lesssim \frac{\sqrt{\pi}}{2}\frac{\Gamma_{\text{as}}}{c_{\text{tot}}}\sqrt{\frac{D_{\max}}{D_{\min}}} = \frac{\sqrt{\pi}}{2}\frac{\Gamma_{\text{as}}}{c_{\text{tot}}}\sqrt{\frac{R_{\max}}{R_{\min}}}, \qquad \text{[S4B]}$$

where $D_{\max}$ and $D_{\min}$ are the diffusion constants of the smallest ($R_{\min}$) and largest ($R_{\max}$) binder in the process, and $c_{\text{tot}}$ is the total concentration of binders. Taking as an example the experimental scenario in Figure 3 ($\Gamma_{\text{as}}$ = 6.8 pmol/cm$^2$, $R_{\text{b-ZZ}}/R_{\text{biotin}} = 5$, and $c_{\text{tot}} = 0.625$ µM), we estimate $\delta_{\text{stagnant,max}} < 100$ µm which is much smaller than the thickness of the well ($H > 1$ mm), as required.

***Worked example for binding with convective fluid transport.*** To model the conditions of QCM-D experiments, we consider a surface with anchor site density $\Gamma_{\text{as}}$ forming one of the walls of a slit of width $2w$, height $2h$, and length $l$. A solution with surface binders flows along the slit with volumetric flow rate $Q$. The inlet of the slit is located at $x = 0$, and the outlet at $x = l$. The binders are homogeneously distributed throughout the solution at the inlet with concentration $c$. In quasi-steady state, the binder surface density is described by [2]

$$\Gamma(x,t) \approx D^{2/3}\left(\frac{3Q}{4h^2wx}\right)^{1/3} ct = \frac{D}{\delta_{\text{flow}}} ct, \qquad \text{[S5]}$$

which gives a depletion layer thickness $\delta_{\text{flow}} = \left(\frac{4Dh^2wx}{3Q}\right)^{1/3}$.

Typical conditions for QCM-D experiments are $2w = 11$ mm, $2h = 0.2$ mm, and the sensing area is centred around $x = 5.5$ mm. The table below provides $\delta$ and $\varepsilon_{\text{flow}}$ (Eq. [S3B]) for the two flow rates used in this study, considering $k_{\text{on}} = 10^7\ \text{M}^{-1}\text{s}^{-1}$, $\Gamma_{\text{as}} = 6.1\ \text{pmol/cm}^2$ (Figure 4D), and $D \leq 580\ \mu\text{m}^2/\text{s}$. It can be seen that the QCM-D flow conditions are within the mass-transport limited regime ($\varepsilon_{\text{flow}} \lesssim 0.01$), and that the depletion layer thickness is nominally inferior to the chamber height ($\delta_{\text{flow}} < 2h$). That we observed deviations from the expected linear dependence in Figure 4 for $Q = 10$ to $20$ µl/min is most likely due to the assumed regular slit not representing the QCM-D chamber geometry very well, leading to a slight underestimation of $\delta_{\text{flow}}$.

| $Q$ | $\delta_{\text{flow}}$ | $\varepsilon_{\text{flow}}$ |
|---|---|---|
| 10 µl/min | <112 µm | <0.0051 |
| 20 µl/min | <89 µm | <0.0064 |
| 200 µl/min | <42 µm | <0.014 |

**2. Numerical analyses of the diffusion and surface binding problem**

We use numerical analyses to quantify the magnitude of the errors made by using Eqs. [4] and [6] in the main text in predicting $\Gamma_{1,\text{sat}}$.

***Binding from stagnant solution.*** We consider a surface with anchor site density $\Gamma_{\text{as}}$ immersed in a semi-infinite solution (Figure S2A). The solution contains two types of surface binders, denoted with subscripts 1 and 2 as in the main text. At the time $t = 0$, the binders are homogeneously distributed throughout the solution at concentrations $c_{1|2}^{(0)}$. The distance from the surface is denoted as $z$. During the binding process, the concentration of binders in the solution $c_{1|2}(z,t)$ and the surface density of binders $\Gamma_{1|2}(t)$ will evolve owing to diffusion of the binders with rate $D_{1|2}$ and binding to the anchor sites with intrinsic rate $k_{\text{on},1|2}$.

To simplify the numerical analysis, we use normalized variables $\tau = \frac{t}{\tau_{\text{transport,stagnant,1}}}$, $u = \frac{zc_1^{(0)}}{\Gamma_{\text{as}}}$, $\hat{\Gamma}_{1|2}(\tau) = \frac{\Gamma_{1,2}(\tau)}{\Gamma_{\text{as}}}$, and $\hat{c}_{1|2}(u,\tau) = \frac{c_{1|2}(u,\tau)}{c_1^{(0)}}$. Using this parametrisation, the time evolution of the system is described by the set of equations

$$\frac{\partial \hat{c}_1}{\partial \tau} = \frac{\partial^2 \hat{c}_1}{\partial u^2}, \quad \text{[S6A]}$$

$$\frac{\partial \hat{c}_2}{\partial \tau} = \frac{D_2}{D_1}\frac{\partial^2 \hat{c}_2}{\partial u^2}, \quad \text{[S6B]}$$

$$\frac{\partial \hat{\Gamma}_1}{\partial \tau} = \begin{cases} \frac{\hat{c}_1(0,\tau)}{\varepsilon_{\text{stagnant,1}}}\left(1 - \hat{\Gamma}_1 - \hat{\Gamma}_2\right) & \text{for } \lambda = 1 \\ \frac{\hat{c}_1(0,\tau)}{\varepsilon_{\text{stagnant,1}}}\left(1 - \frac{\hat{\Gamma}_2}{1-\hat{\Gamma}_1}\right)\left(1 - \lambda\hat{\Gamma}_2\right) & \text{for } \lambda > 1 \end{cases} \quad \text{[S6C]}$$

$$\frac{\partial \hat{\Gamma}_2}{\partial \tau} = \frac{k_{\text{on},2}}{k_{\text{on},1}}\frac{\hat{c}_2(0,\tau)}{\varepsilon_{\text{stll},1}}\left(1 - \hat{\Gamma}_1 - \hat{\Gamma}_2\right), \quad \text{[S6D]}$$

with $\varepsilon_{\text{stagnant,1}}$ defined by Eq. [S3A] for species 1. In Eq. [S6C], the parameter $\lambda$ denotes the number of binding sites occupied by binder 1. Binder 2 is assumed to be small, and so occupies a single binding site. Using these equations, the evolution of the system can be calculated numerically as a function of time. For simplicity, numerical analyses were performed without the numerical pre-factors π/4 in Eq. [S2A] and $4/\pi$ in Eq. [S3A]. Results are shown in Figure S2.

***Binding under flow in a slit.*** To simulate adsorption under convective fluid transport, we use the so-called two-compartment model, which has been shown to accurately describe binding kinetics in transport-limited adsorption [3]. For a single adsorbing species, binding to the surface is assumed to obey, as above, a simplified Langmuir process in which we neglect unbinding, $\partial\Gamma/\partial t = k_{\mathrm{on}} c_{\mathrm{S}}(\Gamma_{\mathrm{as}} - \Gamma)$, where $c_{\mathrm{S}}$ is the near-surface concentration of the binder. The two-compartment model considers two discrete layers in the diffusion and surface-binding problem under flow: a surface layer of thickness $\delta_{\mathrm{S}}$ and binder concentration $c_{\mathrm{S}}$ is located nearest to the surface, and the rest of the depletion layer of thickness $\delta = [4Dh^2wx/(3Q)]^{1/3}$ exhibits a gradient in concentration that goes from the surface value $c_{\mathrm{S}}$ ($c(z \leq \delta_{\mathrm{S}}) = c_{\mathrm{S}}$) to the bulk value $c^{(0)}$ ($c(z = \delta) = c^{(0)}$) (Figure S3A). The time-evolution of $c_{\mathrm{S}}$ is then set by the balance of binder transport towards the surface layer and binder attachment to the surface, yielding

$$\frac{\partial c_{\mathrm{S}}}{\partial t} = \frac{1}{\delta_{\mathrm{S}}}\left[\frac{D}{\delta - \delta_{\mathrm{S}}}(c - c_{\mathrm{S}}) - k_{\mathrm{on}} c_{\mathrm{S}}(\Gamma_{\mathrm{as}} - \Gamma)\right]. \qquad \text{[S7]}$$

We use again normalized variables $\tau = \frac{t}{\tau_{\mathrm{transport,flow,1}}} = \frac{D_1 c_1}{\Gamma_{\mathrm{as}}\delta_1} t$, $\hat{\Gamma}_{1|2}(\tau) = \frac{\Gamma_{1,2}(\tau)}{\Gamma_{\mathrm{as}}}$ and $\hat{c}_{\mathrm{S},1|2} = \frac{c_{\mathrm{S},1|2}}{c_1}$ for two co-adsorbing species. Using this parametrisation, the time evolution of the system is described by the set of equations

$$\frac{\partial \hat{c}_{\mathrm{S},1}}{\partial \tau} = \begin{cases} \frac{\Gamma_{\mathrm{as}}}{C_1\delta_{\mathrm{S}}}\left[1 - \hat{c}_{\mathrm{S},1} - \frac{\hat{c}_{\mathrm{S},1}}{\varepsilon_{\mathrm{flow},1}}\left(1 - \hat{\Gamma}_1 - \hat{\Gamma}_2\right)\right] & \text{for } \lambda = 1 \\ \frac{\Gamma_{\mathrm{as}}}{C_1\delta_{\mathrm{S}}}\left[1 - \hat{c}_{\mathrm{S},1} - \frac{\hat{c}_{\mathrm{S},1}}{\varepsilon_{\mathrm{flow},1}}\left(1 - \frac{\hat{\Gamma}_2}{1-\hat{\Gamma}_1}\right)\left(1 - \lambda\hat{\Gamma}_2\right)\right] & \text{for } \lambda > 1 \end{cases} \qquad \text{[S8A]}$$

$$\frac{\partial \hat{c}_{\mathrm{S},2}}{\partial \tau} = \frac{\Gamma_{\mathrm{as}}}{C_1\delta_{\mathrm{S}}}\left[\frac{D_2(\delta_1 - \delta_{\mathrm{S}})}{D_1(\delta_2 - \delta_{\mathrm{S}})}\left(\frac{c_2}{c_1} - \hat{c}_{\mathrm{S},2}\right) - \frac{k_{\mathrm{on},2}}{k_{\mathrm{on},1}}\frac{\hat{c}_{\mathrm{s},2}}{\varepsilon_{\mathrm{flow},1}}\left(1 - \hat{\Gamma}_1 - \hat{\Gamma}_2\right)\right], \qquad \text{[S8B]}$$

$$\frac{\partial \hat{\Gamma}_1}{\partial \tau} = \begin{cases} \frac{\hat{c}_{\mathrm{S},1}}{\varepsilon_{\mathrm{flow},1}}\left(1 - \hat{\Gamma}_1 - \hat{\Gamma}_2\right) & \text{for } \lambda = 1 \\ \frac{\hat{c}_{\mathrm{S},1}}{\varepsilon_{\mathrm{flow},1}}\left(1 - \frac{\hat{\Gamma}_2}{1-\hat{\Gamma}_1}\right)\left(1 - \lambda\hat{\Gamma}_2\right) & \text{for } \lambda > 1 \end{cases} \qquad \text{[S8C]}$$

$$\frac{\partial \hat{\Gamma}_2}{\partial \tau} = \frac{k_{\mathrm{on},2}}{k_{\mathrm{on},1}}\frac{\hat{c}_{\mathrm{s},2}}{\varepsilon_{\mathrm{flow},1}}\left(1 - \hat{\Gamma}_1 - \hat{\Gamma}_2\right). \qquad \text{[S8D]}$$

with $\varepsilon_{\mathrm{flow},1}$ defined by Eq. (S3B) for species 1, and $\lambda$ as in Eq. [S6C]. Equations [S8A-D] were numerically solved by setting $\delta_{\mathrm{S}} = 2\ \mathrm{nm}$ for both species. Results are essentially insensitive to the exact value of $\delta_{\mathrm{S}}$ as long as $\delta_{\mathrm{S}} \ll \delta$. Results of the numerical analysis, where surface coverages have been evaluated at position *x* = 5 mm, are shown in Figure S3.

**3. Estimation of errors in the control of the surface coverage**

We compare the performance errors of our method with two other possible methods for controlling the grafting density of a functional molecule with an anchor to a surface bearing anchorage sites. The methods are schematically illustrated in Figure S9:

i. In the case of depletion control, the exact number of molecules to graft onto the surface are introduced in the solution (Figure S9A, left).
ii. Kinetic control uses a large amount of material and relies on the incubation time to tune the final surface concentration (Figure S9A, middle).
iii. Competitive binding is the method introduced in this paper (Figure S9A, right).

Here, we consider the case of incubation under static conditions, such as would be used in well plates. The extension to flow conditions is straightforward although the sensitivity to some parameters would be changed.

To estimate the typical errors, we have considered the following parameters as influencing the final grafting density: incubation concentration(s) $c$, incubation time $\tau_{\mathrm{inc}}$, diffusion coefficient(s) $D$, and temperature $T$.

We assume that the different sources of error are independent and thus add up geometrically. As a reference, we considered the case of b-ZZ grafting to a SAv surface in a well of 5 mm diameter filled with 50 µl of solution (a situation representative of a typical well plate), at room temperature.

***Depletion controlled binding.*** In the case of grafting density control through solution depletion, the incubation time is long enough to ensure adsorption of all molecules in solution. The method is thus insensitive to the temperature, the diffusion coefficient and the exact incubation time, but strongly sensitive to the initial concentration in solution $\epsilon_{\text{depletion}} = \delta c/c$. This sensitivity is exacerbated by the small concentration values that are required to functionalize the surface at densities smaller than saturation ($c_{\max} = 0.2$ µg/ml in our example), resulting both in less precision in the exact concentration in solution and a significant influence of non-specific adsorption onto surfaces other than the one to functionalize. This latter effect can account for up to 50% of the proteins in solution for low concentration solutions, and is difficult to mitigate because its amplitude will depend on the environment of the functionalized surface (well, flow chamber, microfluidic circuit, *etc.*) [4]. Here we have considered an induced uncertainty of 20% on the effective concentration of reagent in solution. Assuming a minimum pipetting error of at least 0.5 µl or 5% of the pipetted volume, 2 dilution steps (the second depending on the final required concentration $\gamma c_{\max}$), and independent addition of errors due to pipetting and non-specific adsorption, we obtain $\epsilon_{\text{depletion}} = \sqrt{\left(\frac{\delta c}{c}\right)^2_{\text{dilution 1}} + \left(\frac{\delta c}{c}\right)^2_{\text{dilution 2}} + \left(\frac{\delta c}{c}\right)^2_{\text{depletion}}} = \sqrt{0.05^2 + \left(\frac{0.0025}{\gamma}\right)^2 + 0.20^2}$, with $\gamma$ the relative surface coverage ($\gamma = 1$ corresponding to saturation). In this formula, we have assumed that the functionalized area and the incubation volume are known perfectly as the resulting errors are expected to be of much smaller amplitude.

***Kinetically controlled binding.*** In the case of kinetic control, the grafting density is tuned through the incubation time of a solution of known concentration, following Eq. [1] in the main text. To estimate the relative errors in time and concentration, we assume that the incubation time for saturation was set to 2 h and calculated the resulting concentration, yielding a concentration of 0.5 µg/ml which can typically be achieved with a 10% error. The incubation time, in turn, is assumed to be controlled with a 1 min precision which accounts for the time required for mixing the solution at the onset of incubation, and the successive rinsing steps at the end of the process. Since the incubation time scales with the square of the surface concentration, this yields an error $\frac{1}{2}\frac{\delta\tau_{\text{inc}}}{\tau_{\text{inc}}} = \frac{1\text{ min}}{2 \times 120\text{ min} \times \gamma^2} = \frac{1}{240\,\gamma^2}$. In addition, temperature plays a critical role in this situation as it influences the diffusion coefficient of the molecule directly, and through changes in the liquid viscosity (assumed to be close to that of water). Here we assumed that the ambient temperature could change by up to 5°C, yielding an associated 6% error. This source of variability can be mitigated by working in a temperature-controlled environment, but is actually not a main source of error here. Finally, we assume the hydrodynamic radius of the molecule to be known with 10% precision. Combining the different abovementioned independent sources of error, we obtain $\epsilon_{\text{kinetic}} = \sqrt{\left(\frac{\delta c}{c}\right)^2 + \left(\frac{1}{2}\frac{\delta\tau_{\text{inc}}}{\tau_{\text{inc}}}\right)^2 + \left[\left(1 + \frac{T}{\eta}\frac{d\eta}{dT}\right)\frac{\delta T}{T}\right]^2 + \left(\frac{\delta r_1}{r_1}\right)^2}$ with $\eta$ the solution viscosity, which gives $\epsilon_{\text{kinetic}} = \sqrt{0.10^2 + \left(\frac{1}{240\,\gamma^2}\right)^2 + 0.06^2 + 0.10^2}$.

***Competitive binding.*** In the case of the control of grafting density through mixing with a free anchor, the relevant uncertainties relate to the concentrations and hydrodynamic radii of the two species. The relative errors in these quantities typically scale by $\sqrt{2}$. However, the concentrations used can be much higher than for the first two methods, and thus are supposed to be known with 5% accuracy. As above, the error on the hydrodynamic radius is estimated to be 10% for each molecule. In addition, the method is insensitive to the temperature (as only the ratio of the diffusion constants appears in Eq. [4] in the main text) and to the incubation time (as saturation of the anchorage sites is reached). For the sake of completeness, we also consider the case where not only a relative fraction of occupancy of the anchorage sites is needed, but an absolute value for surface coverage: with the mixing method this will add an uncertainty due to the imperfect knowledge of the number of anchorage sites. In the case of a monolayer of SAv attached to a SLB, the uncertainty (combining the density of SAv and the number of available biotin binding sites per SAv [5]) is estimated to be around 10%. We thus obtain the error estimates $\epsilon_{\mathrm{cb,fractional\ occupancy}} = \sqrt{(1-\gamma)^2\left[\left(\frac{\delta c_1}{c_1}\right)^2 + \left(\frac{\delta c_2}{c_2}\right)^2 + \left(\frac{1}{2}\frac{\delta r_1}{r_1}\right)^2 + \left(\frac{1}{2}\frac{\delta r_2}{r_2}\right)^2\right]} = \sqrt{2(1-\gamma)^2\left(0.05^2 + \frac{0.10^2}{4}\right)}$, and $\epsilon_{\mathrm{cb,surface\ density}} = \sqrt{(1-\gamma)^2\left(\left(\frac{\delta c_1}{c_1}\right)^2 + \left(\frac{\delta c_2}{c_2}\right)^2 + \left(\frac{1}{2}\frac{\delta r_1}{r_1}\right)^2 + \left(\frac{1}{2}\frac{\delta r_2}{r_2}\right)^2\right) + \left(\frac{\delta \Gamma_{as}}{\Gamma_{as}}\right)^2} = \sqrt{2(1-\gamma)^2\left(0.05^2 + \frac{0.10^2}{4}\right) + 0.10^2}$.

Figure S9B shows the resulting errors as a function of the expected surface coverage, demonstrating that the proposed approach is expected to be more precise over a wide range of surface coverages.

**SUPPORTING TABLE**

**Table S1.** Masses, hydrodynamic radii and diffusion coefficients of molecules with biotin anchor used in this study.

| **Molecule** | ***M*$_\mathrm{W}$** | **Hydrodynamic radius** | **Diffusion coefficient** |
|---|---|---|---|
| biotin | 244.3 Da | $R_\mathrm{biotin} = 0.37\ \mathrm{nm}$<br>From van der Waals volume ( $V_\mathrm{vdW} = 0.213\ \mathrm{nm}^3$ ) assuming a sphere, as previously validated by others [6]. | $D_\mathrm{biotin} = 580\ \mathrm{\mu m^2/s}$<br>From the hydrodynamic radius [a)]. |
| b-FITC | 732.8 Da | $R_\mathrm{b-FITC} = 0.63\ \mathrm{nm}$<br>From the diffusion coefficient [a)]. | $D_\mathrm{b-FITC} = 340\ \mathrm{\mu m^2/s}$<br>As previously determined by others [7]. |
| b-ZZ | 16.2 kDa | $R_\mathrm{b-ZZ} = 2.0\ \mathrm{nm}$<br>From the diffusion coefficient [a)]. | $D_\mathrm{b-ZZ} = 107\ \mathrm{\mu m^2/s}$<br>Computed following Ref. [8] using a radius of gyration $R_\mathrm{g} = 1.05 \times \left(\frac{3}{5}\right)^{1/2} \left(\frac{3}{4\pi}\frac{M_\mathrm{W}}{\rho N_\mathrm{A}}\right)^{1/3} = 1.4\ \mathrm{nm}$ , with density $\rho = 1.4\ \mathrm{g/cm^3}$ (typical for proteins); the factor 1.05 accounts for the biotin tag likely pointing out of the protein. |
| alkoxyamine-EG$_4$-biotin | 434.2 Da | $R_\mathrm{b-alkoxyamine} = 0.6\ \mathrm{nm}$<br>From the hydrodynamic radius of the EG$_4$ spacer (0.4 nm) [9], estimating that the biotin and alkoxyamine moieties add a further 0.2 nm. | $D_\mathrm{biotin} = 360\ \mathrm{\mu m^2/s}$<br>From the hydrodynamic radius [a)]. |
| GAG-b (biotinylated CS-D with $n_\mathrm{ds} = 76..90$ disaccharides | 38..45 kDa | $R_\mathrm{GAG-b} \approx 7..8\ \mathrm{nm}$<br>$R_\mathrm{GAG-b} \approx 0.50\ \mathrm{nm}\ (n_\mathrm{ds})^{0.61}$ , with $n_\mathrm{ds}$ the number of disaccharides, following analysis by Ref. [10] of the $M_\mathrm{W}$ dependence of the hydrodynamic radius of hyaluronan (HA) in 0.2 M NaCl. We estimate that sulfated GAGs such as CS have a similar hydrodynamic radius at 150 mM NaCl and comparable $n_\mathrm{ds}$. The disaccharide mass is $M_\mathrm{W,ds}$ = 0.4 kDa for HA, and estimated at 0.5 kDa for sulfated GAGs. | $D_\mathrm{GAG-b} \approx 28..31\ \mathrm{\mu m^2/s}$<br>From the hydrodynamic radius [a)]. |

a) Hydrodynamic radii and diffusion coefficients were interconverted using the Stokes Einstein relation $D = k_\mathrm{B}T/(6\pi\eta R)$, with solvent viscosity $\eta = 1.0\ \mathrm{mPa\ s}$ and thermal energy $k_\mathrm{B}T$ at $T$ = 20 °C.

**SUPPORTING FIGURES**

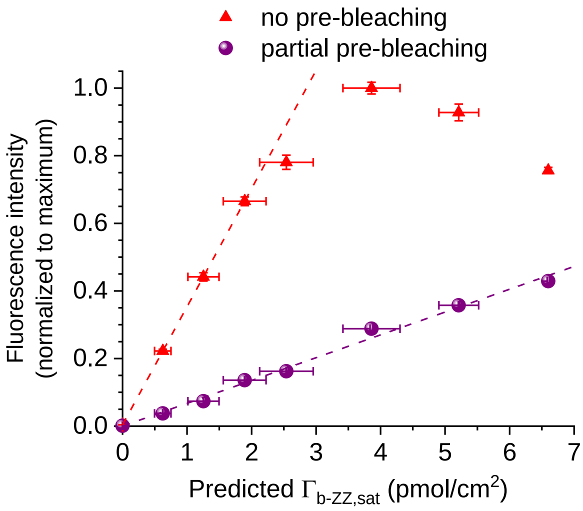


**Figure S1. Partial pre-bleaching of b-FITC to eliminate self-quenching on the surface.** The b-FITC in solution was pre-bleached before surface functionalisation using a halogen lamp (KL 1500 LCD; Schott, Germany) covering the entire visible spectrum and hence exciting and bleaching the FITC fluorophore efficiently. The fluorophore solution (20 µg/ml in working buffer) was installed in a transparent glass vial at a distance of 1.5 cm from the light source and illuminated. During the illumination process, the solution was briefly stirred on the vortex once per minute to maintain a homogeneous solution. Fluorescence emission spectra were determined with a spectrofluorometer (FluoroMax-4; Horiba, Japan) in plastic cuvettes (UV-Cuvette micro 70 µl; BRAND GmbH, Germany) before illumination and after 30 min of illumination. The graph shows the fluorescence intensity measured by confocal microscopy for surfaces functionalized with different b-FITC/biotin mixes; the predicted surface coverage was determined through Eq. [4] with $\Gamma_{as} = 6.8\ \mathrm{pmol/cm^2}$. For non-bleached fluorophores (red triangles), the intensity of fluorescence is linearly increasing with the surface density of b-FITC until an intermolecular distance of 8 nm (b-FITC surface density: 2-2.5 pmol/cm$^2$), reaching a maximum for an intermolecular distance of 6 nm (b-FITC surface density: 4-5 pmol/cm$^2$), and then decreasing due to intermolecular quenching. In contrast, the dependence is linear across the full range of surface coverages for the b-FITC that was illuminated for 30 min (purple spheres; same data as shown in Figure 2B), indicating that the illumination induced sufficient bleaching to avoid self-quenching of surface-anchored b-FITC.

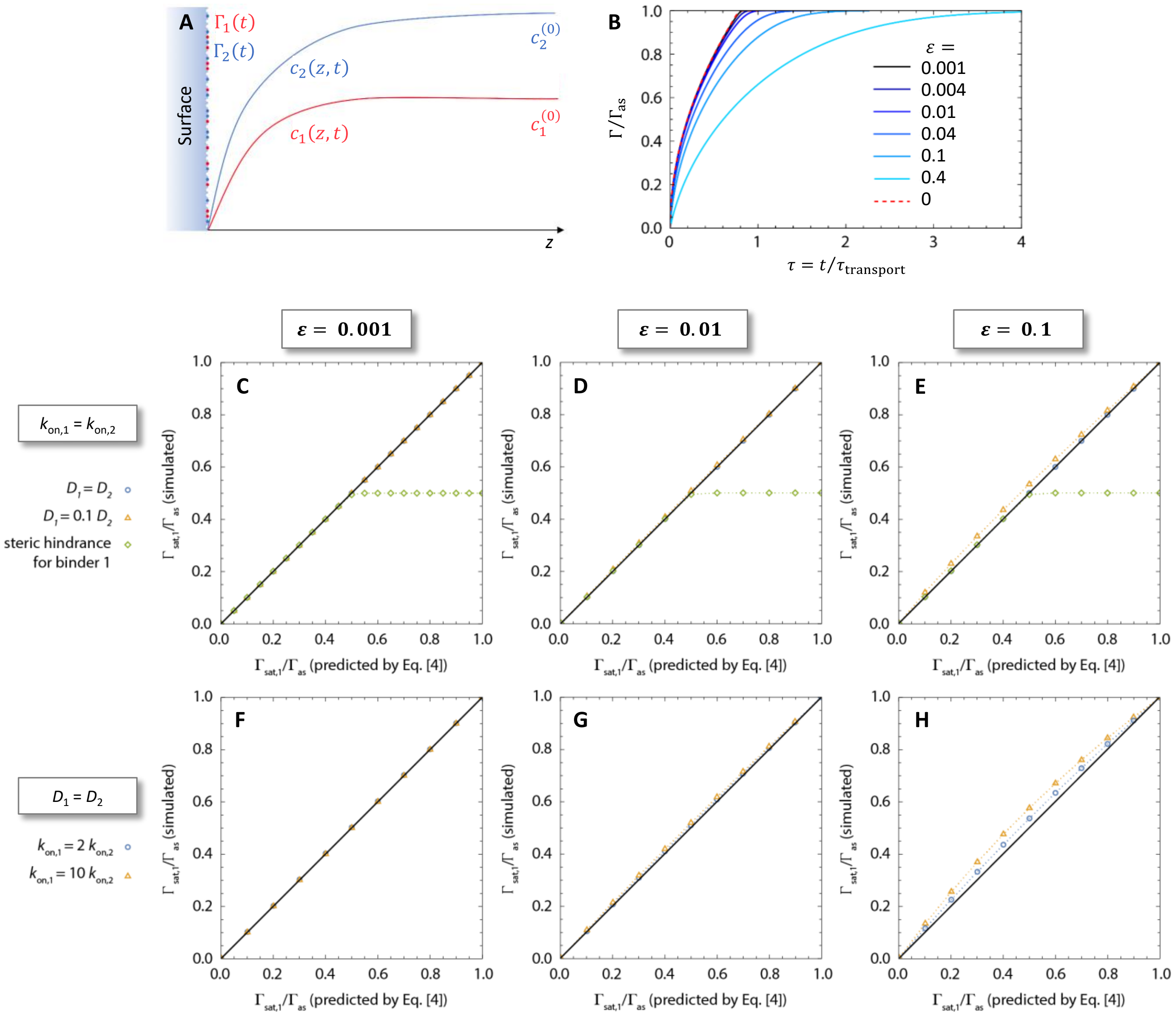


**Figure S2. Numerical analysis of the diffusion and surface binding problem in stagnant solution. A.** Schematic of the binding scenario. **B.** Kinetics of surface binding for a single binder at different bulk concentrations, defined through $\varepsilon_{stagnant} = cD/(k_{on}\Gamma_{as}^2)$ (with color codes as indicated). The perfect sink condition (Eq. [1]; dashed red line) is equivalent to $\varepsilon_{stagnant} \to 0$, and predicts saturation ($\Gamma/\Gamma_{as} = 1$) at $\tau = \pi/4 \approx 0.8$. At $\varepsilon_{stagnant} = 0.1$, the time to 99% saturation approximately doubles. **C-F.** Tests of the prediction accuracy of Eq. [4], with differences in $D_{1|2}$ or $k_{on,1|2}$ as indicated (blue circles and orange triangles). (C-D, F-G) For $\varepsilon_{stagnant,1} \leq 0.01$, neither a 10-fold differences in $D$ nor a 10-fold difference in $k_{on}$ impact the predictions appreciably. (E, H) For $\varepsilon_{stagnant} = 0.1$, a 10-fold difference in $D$, or a 2-fold difference in $k_{on}$, entail up to 10% deviations. Steric hindrance (C-E; green lozenges - each binder 1 molecule occupies $\lambda = 2$ anchor sites) does not impact the predictions until close to saturation at $\Gamma_{sat,1}/\Gamma_{as} = 1/\lambda$, followed by a plateau.

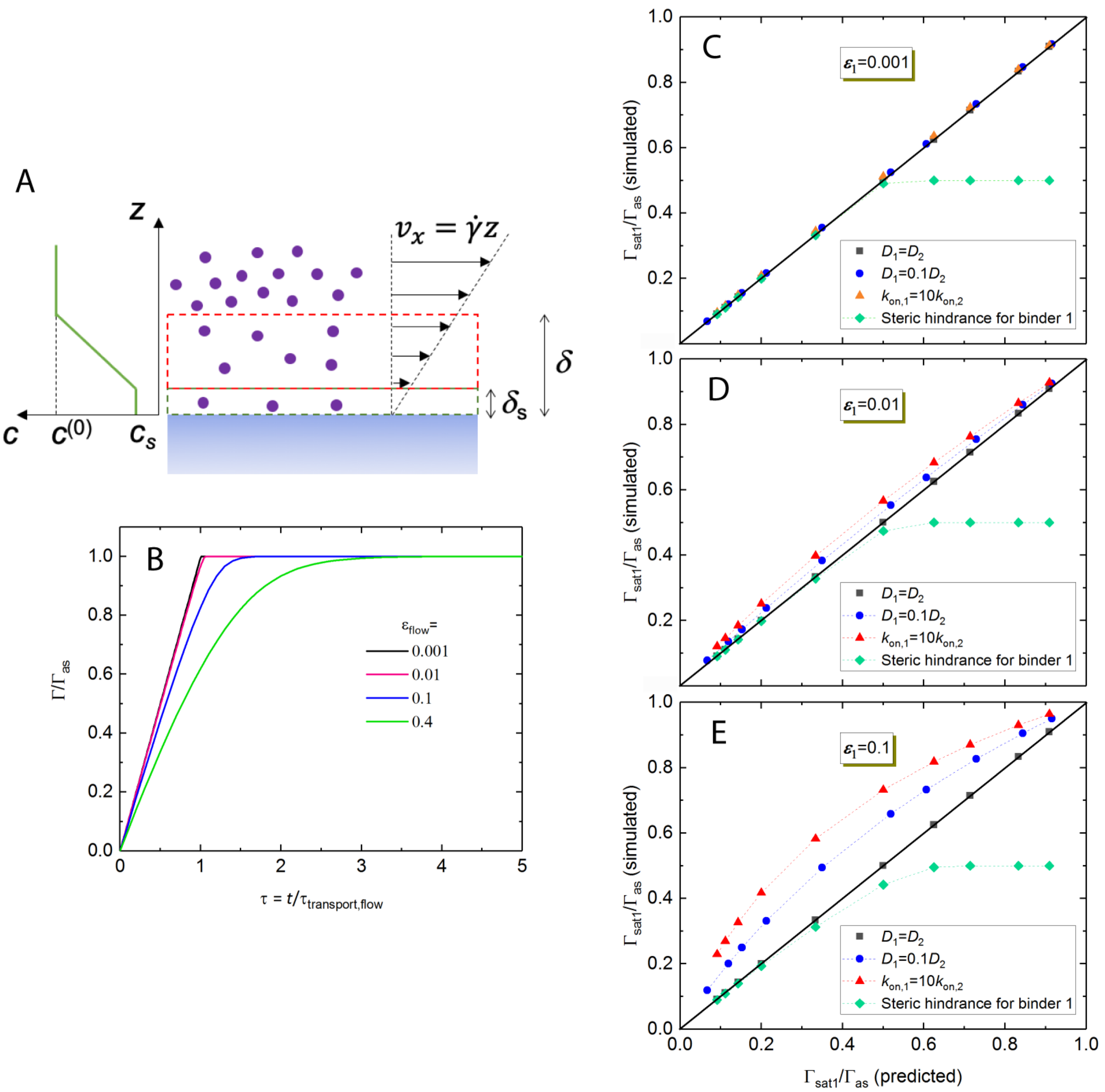


**Figure S3. Numerical analysis of the transport and surface binding problem under flow. A.** Sketch of the two-compartment model. The depletion layer with concentration gradient is highlighted by a dashed red rectangle. The shear rate at the wall is defined as $\dot{\gamma} = \frac{3Q}{4h^2 w}$ for a flow rate $Q$ in a rectangular channel of height $2h$ and width $2w$. **B.** Kinetics of surface binding for a single binder at different $\varepsilon_{\text{flow}} = D/(k_{\text{on}}\Gamma_{\text{as}}\delta)$ (with color codes as indicated). For $\varepsilon_{\text{flow}} = 0.001$, the binding rate is essentially constant (mass transport limited) up to saturation and saturation is reached at $\tau = 1$, equivalent to the perfect sink condition. At $\varepsilon_{\text{flow}} = 0.1$, the time to 99% saturation approximately doubles, and deviations from the perfect sink condition become noticeable at 20% saturation. **C-E.** Tests of the prediction accuracy of Eq. [6], with differences in $D_{1|2}$ or $k_{\text{on},1|2}$ as indicated. (C) For $\varepsilon_{\text{flow},1} = 0.001$, neither a 10-fold differences in $D$ nor a 10-fold difference in $k_{\text{on}}$ impact the predictions appreciably. Steric hindrance (green lozenges - each binder 1 molecule occupies $\lambda = 2$ anchor sites) does not impact the predictions until close to saturation at $\Gamma_{\text{sat},1}/\Gamma_{\text{as}} = 1/\lambda$, followed by a plateau. (D) For $\varepsilon_{\text{flow},1} = 0.01$, a 10-fold difference in $D$ entails up to 15% deviations, and a 10-fold difference in $k_{\text{on}}$ up to 25% deviations. (E) For $\varepsilon_{\text{flow},1} = 0.1$, a 10-fold difference in $D$ entails up to 50% deviations, and a 10-fold difference in $k_{\text{on}}$ up to 60% deviations.

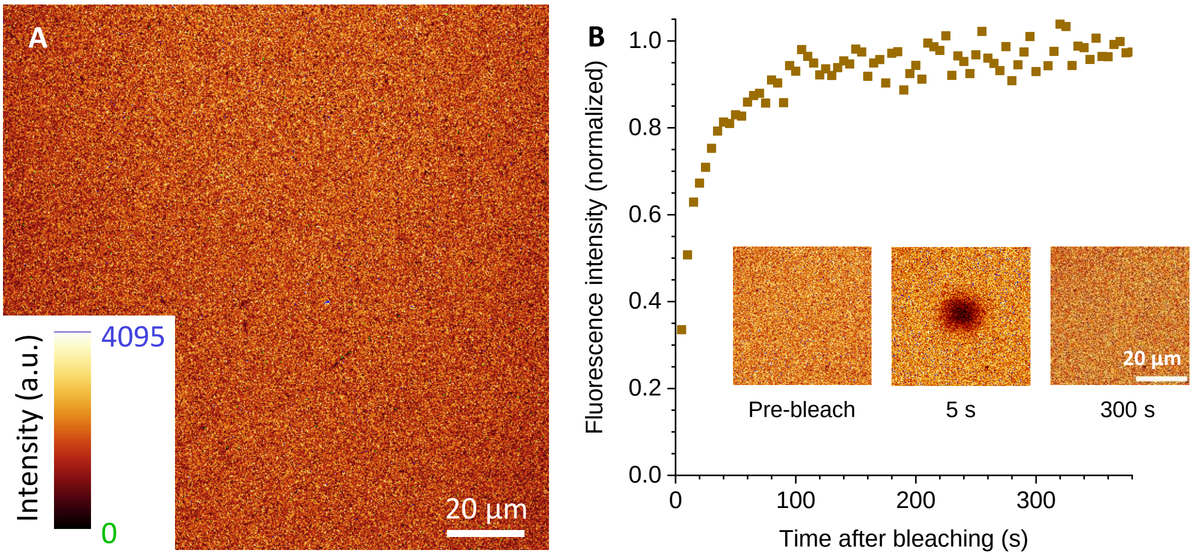


**Figure S4. Fluorescence analysis of the streptavidin monolayer on a glass-supported lipid bilayer. A.** Representative micrograph with fluorescently labelled SAv, demonstrating that SAv coverage is essentially uniform. A few small defects are noticeable as darker patches with a typical size of 1 µm or less, but these occupied only a small fraction of the total surface area and negligibly impacted the data analysis. **B.** Fluorescence recovery after photo-bleaching demonstrates in-plane mobility of SAv on the SLB. A square of 10 µm width in the centre of the image was photo-bleached, and fluorescence recovery following in-plane diffusion of SAv was then monitored (insets show snapshots of the bleached area, with fluorescence intensities normalized against unbleached areas). The intensity in the bleached area essentially returned to the level of unbleached reference areas, indicating that >95% of the SAv molecules are in-plane mobile on the fluid SLB. Conditions: see Materials and Methods - Surface preparation for confocal microscopy.

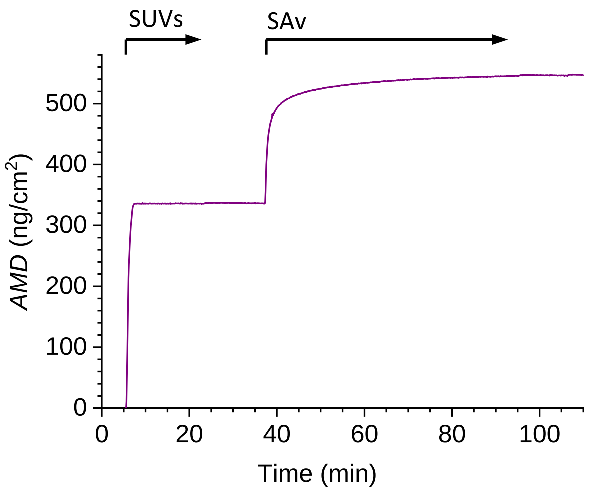


**Figure S5. Representative SE binding assay for the formation of SLBs and SAv monolayers.** Shown is the areal mass density, *AMD*, determined from time-resolved SE analysis (with refractive index increments d*n*/d*c* = 0.169 $cm^3$/g for lipids, and 0.180 $cm^3$/g for SAv). The start and duration of the incubation with different samples are indicated by solid arrows atop the graph. There are rinsing steps with working buffer before and after each sample incubation step. The areal mass density of SAv at the end of the incubation process was 211 ± 9 ng/$cm^2$ (mean ± standard deviation) across all surface preparations ($n$ = 8), illustrating good reproducibility of the surface preparation. This areal mass density corresponds to a molar surface density of 3.52 ± 0.15 pmol/$cm^2$. Conditions: SUVs – 50 µg/ml, containing 5 mol-% DOPE-CAP-biotin in a DOPC background; SAv – 0.33 µM.

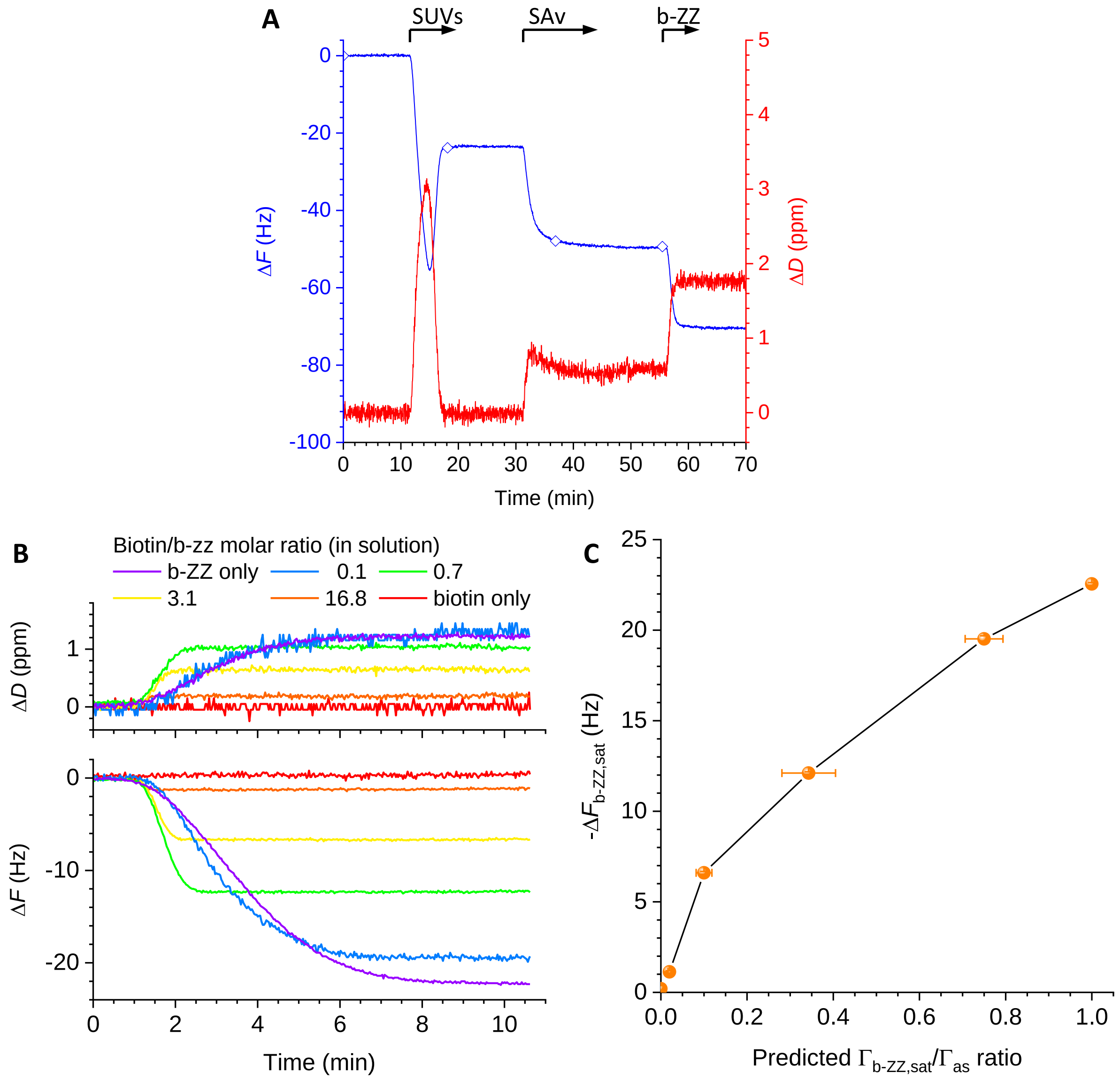


**Figure S6. Representative QCM-D binding assay for the formation of SLBs and a SAv monolayer, followed by b-ZZ grafting. A.** Shown are the QCM-D frequency shifts ($\Delta F$ – blue line with diamond symbols) and dissipation shifts ($\Delta D$ – red line) for the 5$^{th}$ overtone ($i$ = 5) as a function of time. The start and duration of the incubation with different samples are indicated by solid arrows atop the graph. There are rinsing steps with working buffer before and after each sample incubation step. The two-phase response upon b-SUV incubation is characteristic for SUVs initially binding intact and then rupturing and spreading as the surface coverage increases; the final shifts upon SUV incubation ($\Delta F$ = -24 ± 1 Hz and $\Delta D$ < 0.2 ppm) indicate an SLB of good quality (*i.e.*, with minimal residual liposomes) has formed [11]. The final shifts upon SAv incubation ($\Delta F$ = -26 ± 1 Hz and $\Delta D$ = 0.5 ppm) are fully consistent with the formation of a SAv monolayer [5, 12]. b-ZZ incubation demonstrated rapid binding of the protein (within 1-2 minutes) until full saturation, with final shifts $\Delta F$ = -21 ± 1 Hz and $\Delta D$ = 1.2 ± 0.1 ppm. The b-ZZ layer remained stable upon rinsing in buffer. Conditions: SUVs – 50 µg/ml, containing 5 mol-% DOPE-CAP-biotin in a DOPC background; SAv – 0.33 µM, b-ZZ – 14 µM. **B-C.** Data equivalent to Figures 4A-B, but in a reduced flow rate regime of 10 µl/min (biotin/b-ZZ molar ratio of 0.1, as well as b-ZZ only and biotin only) to 20 µl/min (biotin/b-ZZ molar ratios of 0.7, 3.1 and 16.8).

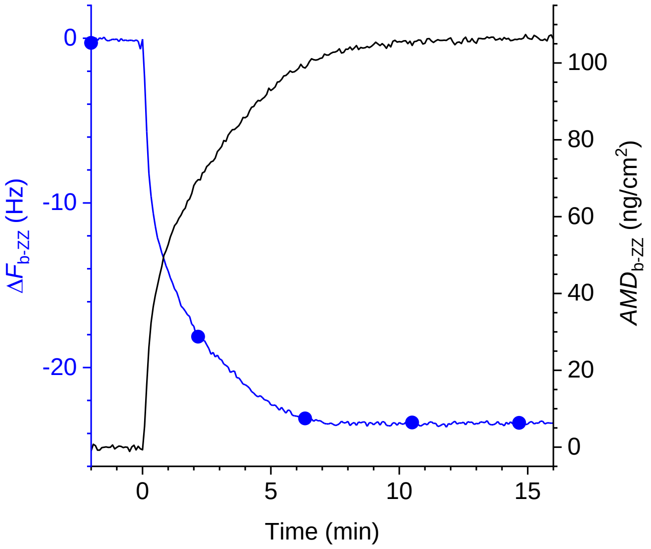


**Figure S7. Establishing a standard curve linking QCM-D frequency shifts to b-ZZ surface coverage on a SAv-on-SLB surface.** Shown are the frequency shift ($\Delta F_{\mathrm{b-ZZ}}$ – blue line with circles; determined by QCM-D, *i* = 5) and the areal mass density ($AMD_{\mathrm{b-ZZ}}$ - black line; determined by SE) acquired simultaneously on a QCM-D sensor surface as a function of time for the binding of b-ZZ to a SAv-coated SLB. The standard curve in Figure 4C was obtained by correlating $\Delta F_{\mathrm{b-ZZ}}$ and $\Gamma_{\mathrm{b-ZZ}} = AMD_{\mathrm{b-ZZ}}/M_{\mathrm{W,b-ZZ}}$. Conditions: b-ZZ – 0.31 µM, injected at 0 min; incubation proceeded in stagnant solution, except for the first 10 s when the solution was stirred to homogeneity. Surface preparation (not shown, but analogous to what is shown in Figures S4 and S5): SUVs - 50 µg/ml, containing 5 mol-% DOPE-CAP-biotin in a DOPC background, were incubated under stirring for 7 min until SLB formation was complete; SAv – 0.13 µM, was then incubated for 30 min (under stirring for 20 min, followed by 10 min in stagnant solution). Data were acquired with an open fluid cell, as described in detail in Ref. [13]; a QSX303 sensor was deployed as a substrate, and SE data were fitted to extract the *AMD* as described in Ref. [14].

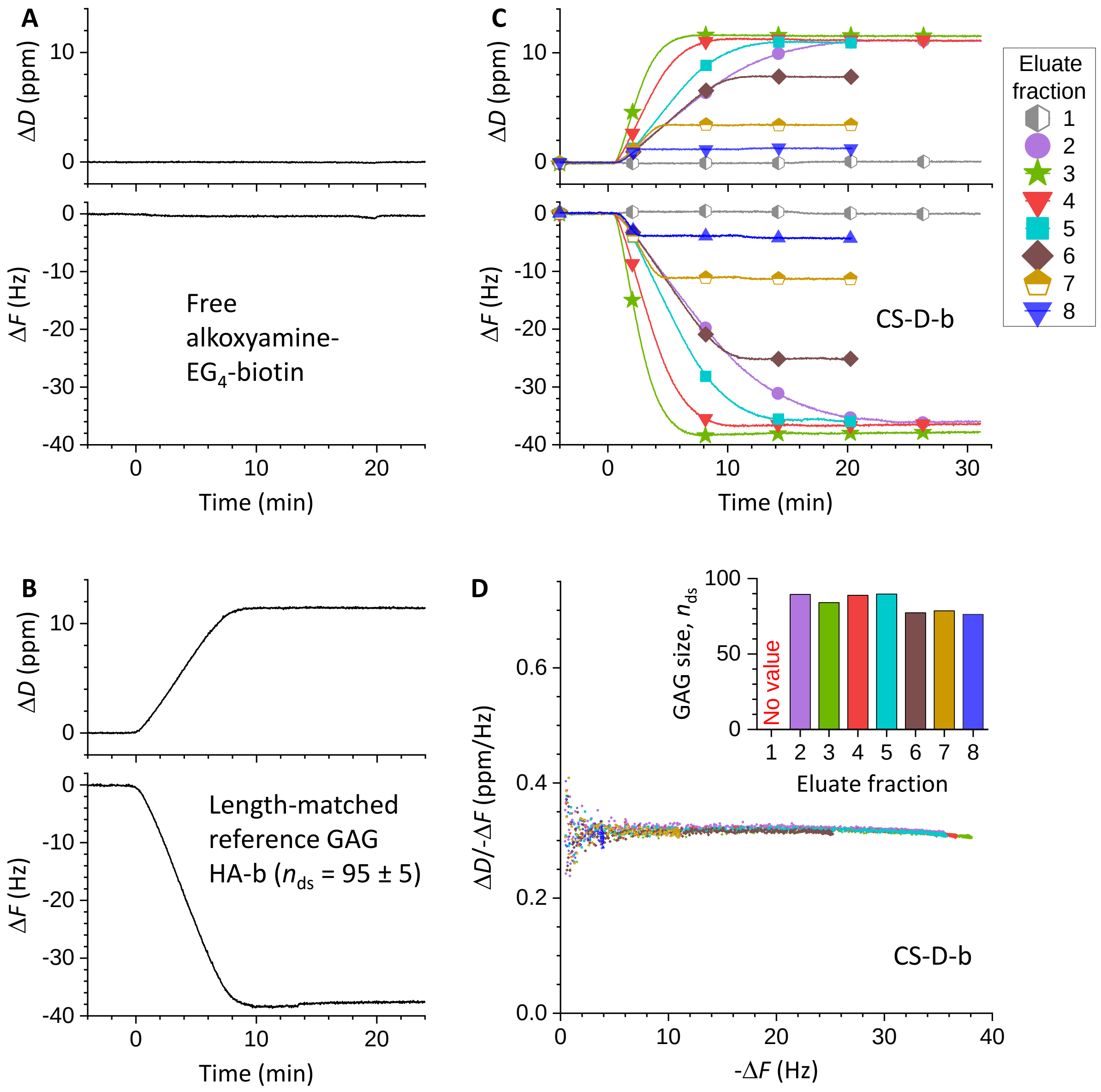


**Figure S8. Complementary data for post-chromatographic QCM-D analysis of biotinylated CS-D GAG samples**. **A.** QCM-D responses ($\Delta F$ - bottom; $\Delta D$ - top; $i$ = 5) for the binding of free alkoxyamine-EG$_4$-biotin. Conditions: 1 µM alkoxyamine-EG$_4$-biotin were incubated from 0 to 20 min; working buffer was flown over the sensor for the remaining times; flow rate – 20 µL/min. **B.** QCM-D responses ($\Delta F$ - bottom; $\Delta D$ - top; $i$ = 5) for the binding of biotinylated hyaluronan (HA-b) of 38 ± 2 kDa (95 ± 5 disaccharides), which was used as a reference. Conditions: HA-b was incubated from 0 to 10 min; working buffer was flown over the sensor for the remaining times; flow rate – $Q_{\mathrm{ref}}$ = 10 µl/min; HA-b concentration – $c_{\mathrm{ref}} = 0.13$ µM. **C.** QCM-D responses ($\Delta F$ - bottom; $\Delta D$ - top; $i$ = 5) for the binding of biotinylated CS-D GAG (CS-D-b), for each of the eluate fractions (EFs) retrieved from the size-exclusion column (with symbols and colours as indicated). Conditions: CS-D-b incubation started at 0 min, and proceeded for 16 min, except for EF2, which proceeded for 24 min (during other times, plain working buffer was flown over the sensor); flow rate – $Q_{\mathrm{sample}}$ = 20 µl/min; CS-D-b concentration – the EFs as retrieved from the size-exclusion column were diluted 15-fold for QCM-D analysis. Max rate, -$\Delta F_{\text{GAG-b}}/\Delta t$, and Max response, -$\Delta F_{\text{GAG-b,sat}}$, were extracted for each EF, and are reported in Figures 6C and 6E, respectively. CS-D-b concentrations (Figure 6D) were calculated from the max rates as $c_{\mathrm{sample}} = f_{\mathrm{dilution}} c_{\mathrm{ref}} \frac{\Delta F_{\mathrm{sample}}/\Delta t}{\Delta F_{\mathrm{ref}}/\Delta t} \left(\frac{Q_{\mathrm{ref}}}{Q_{\mathrm{sample}}}\right)^{1/3}$, with the dilution factor $f_{\mathrm{dilution}} = 15$. The index 'sample' here refers to any of the EFs, and the index 'ref' to the length-matched HA-b reference sample shown in panel B ($\Delta F_{\mathrm{ref}}/\Delta t$ = 5.4 Hz/min). **D.** Parametric plots of $\Delta D$/-$\Delta F$ versus -$\Delta F$ (here used as a proxy for GAG surface coverage)

during the GAG-b film formation. The parametric plots virtually superpose for the 7 eluate fractions with GAG content, demonstrating that the size exclusion chromatography, whilst effective in separating free biotin from the GAGs, hardly separated the pool of CS-D molecules by their size. The $\Delta D/\text{-}\Delta F$ values at $-\Delta F$ = 2.5 Hz were used to determine the effective GAG-b size (in number of disaccharides, $n_{ds}$; following the procedure described in Ref. [15]) in each fraction (inset). This analysis confirmed that the effective GAG size was essentially unchanged across fraction 2 to 5 (with approximately 90 disaccharides per chain on average), and only marginally reduced (to approximately 80 disaccharides) for the later fractions, and that the HA-b reference sample shown in panel B was approximately length matched as desired.

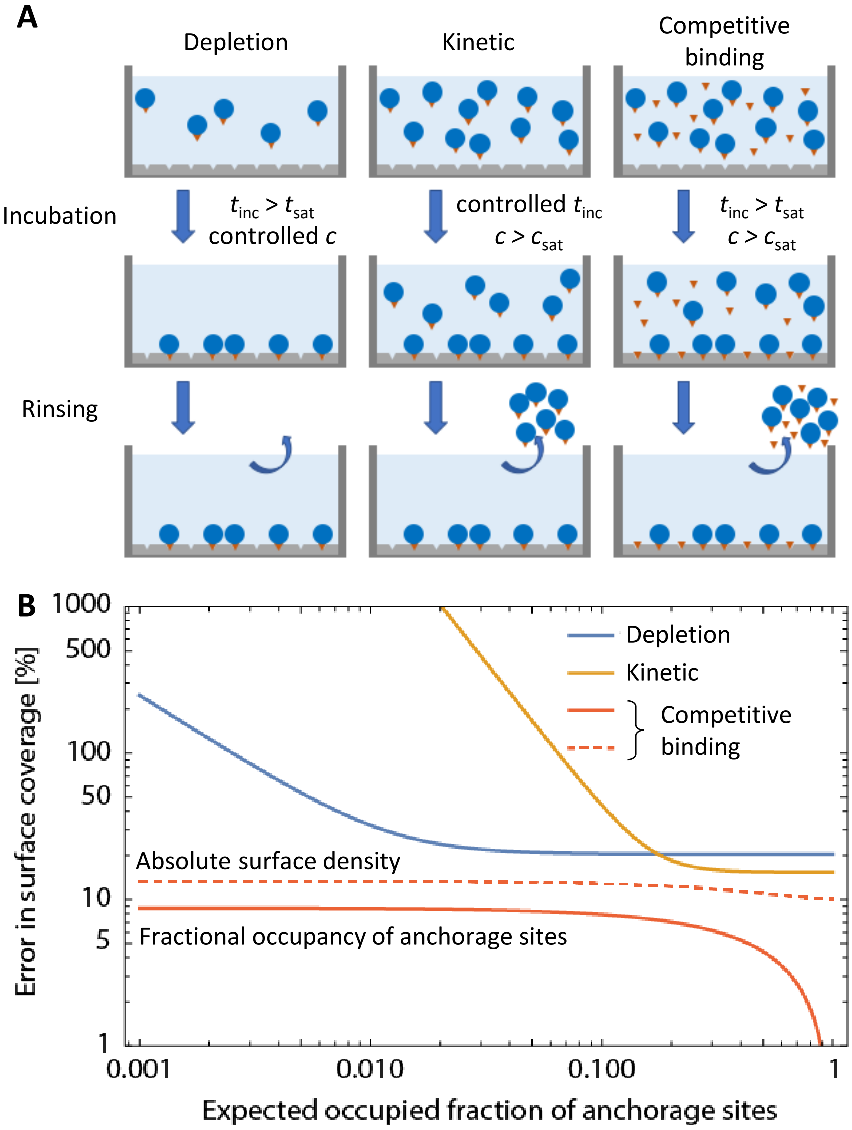


**Figure S9. Comparative error analysis for methods to control molecular surface densities. A.** Illustrations of the considered methods for controlling the grafting density of a functional molecule with anchor to a surface with anchorage sites ($c_{\text{sat}}$ – saturating concentration(s); $t_{\text{inc}}$ – incubation time; $t_{\text{sat}}$ – time required for surface saturation or solution depletion). **B.** Relative errors in surface coverage as a function of the fraction of occupied anchorage sites, for the 3 methods described in panel A.
